# Calcium Binding Affinity in the Mutational Landscape of Troponin-C: Free Energy Calculation, Coevolution Modeling and Machine Learning

**DOI:** 10.1101/2024.09.09.612070

**Authors:** Pooja, Pradipta Bandyopadhyay

## Abstract

Mutation in calcium-binding proteins (CBPs) can significantly influence *Ca*^2+^ binding affinity (BA), resulting in substantial impairment in the signaling process and leading to several lethal diseases. The knowledge behind the changes in the binding affinity can help in understanding the signaling process and designing inhibitors for therapeutic usage. However, accurate prediction of BA for a large number of mutations has been elusive. In this work, for an important calcium binding protein, cardiac Troponin-C, we have developed an integrative modeling approach that combines molecular dynamics (MD)-based binding free energy calculations, prediction of plausible mutants using evolutionary information, and an interpretable machine learning model to predict *Ca*^2+^ BA for a large number of mutations (seventy-six in all). For the binding free energy calculation, we have used a charge-scaling based MD simulation that considers the polarization in the system, which is critical for divalent ion binding with proteins. The well-known molecular mechanics Poisson-Boltzmann surface area (MM-PBSA) method was used for the binding free energy calculations. The calculated results for twenty-four disease mutants, which are associated with different cardiomyopathies and have experimental binding affinity, are in close agreement with the experimental results. To study other plausible mutations, we have probed the evolutionary landscape of cardiac Troponin-C and used the EVmutation method of Hopf *et al.*(Nature biotechnology 2017, 35, 128–135) to generate sixty-one additional mutants. Finally, a Support vector regression model was developed for both observed and plausible mutations. Our machine learning model used simple structure and sequence-based descriptors along with MD-based descriptors and gave a mean squared error (MSE) of only 0.16 kcal/mol. Assessment of the contribution of each descriptor shows that the number of water molecules within the *Ca*^2+^ binding site, type of amino acid substitution (e.g. polar to hydrophobic reduces the binding affinity), and the distance of mutation with *Ca*^2+^ are the most important factors in determining the binding affinity. This integrative modeling can be used for other CBPs and can lay the path for modeling the complex and astronomically large mutational landscape of Calcium-binding proteins.

## 1. Introduction

Calcium-binding proteins (CBPs) are a diverse group of intracellular signaling molecules characterized by their ability to bind calcium ions through specific domain. These proteins act as calcium sensors inside the cell and mediators of various signaling pathways that regulate cellular processes related to muscle contraction, neuronal signal transduction, and cell proliferation.^1,2^ When calcium ion binds to CBPs such as troponin-C, calmodulin, and paralbumin, these proteins undergo conformational changes resulting in the exposure of a hydrophobic patch, which interacts with the target protein and cellular process occurs. ^3^ The binding of *Ca*^2+^ in the conserved helix-loop-helix motif (EF-hand loop) of these proteins, is facilitated by the presence of specific coordinating residues in a pentagonal bipyramidal geometry.^4^ Even a point mutation in the critical amino acid residue of these proteins disrupts *Ca*^2+^ binding affinity, affecting its functionality, and may lead to severe diseases including cardiomyopathies, neurodegenerative disorders, and cancer.^5–7^ Due to the profound significance of these proteins in the regulation of cellular pathways, it has been the subject of thorough investigation, encompassing both experimental studies using Isothermal Titration Calorimetry, Fluorescence spectroscopy, and Surface Plasmon Resonance^8–13^ and theoretical^7,14–19^ approaches. However, a complete understanding of the mechanism that governs the *Ca*^2+^ binding properties of CBPs still eludes us. Accurate estimation of *Ca*^2+^ binding affinity to the different variants (wild-type and mutants) of CBPs is extremely important as alteration of the binding affinity for mutants can have potentially severe consequences. For instance, mutation in cardiac troponin-C (cTn-C) protein changes the *Ca*^2+^ binding affinity towards protein variant, which can lead to cardiac diseases such as hypercardiomyopathy and dilated or restricted cardiomyopathy.^6^ Likewise, mutation in calmodulin (CaM) protein affects *Ca*^2+^ binding affinity and thus affects the regulatory pathways that may cause cardiac arrhythmia, long QT syndrome (LQTS), and Alzheimer’s disease.^20,21^ Experimental studies on determining the binding affinity have been extremely valuable; however, these are not possible to conduct for the large mutational space of hundreds of CBPs in mammalian systems, although techniques like deep mutational scan^22^ may circumvent this to some extent. Hence, the role of an accurate computational prediction method is immensely important. Thus, the need of the hour is to develop a significantly accurate and efficient prediction model to estimate the *Ca*^2+^ binding affinity using simple descriptors of protein variants. Understanding the reasons for the change in binding affinities can give avenues to modulation of *Ca*^2+^ binding and eventually to CBP design. For the calculation of *Ca*^2+^ binding affinity to CBPs, various structure, and sequence-based approaches have been used sporadically, but still, there are several technical challenges.^19,23–25^ For instance, work done by Gourinath’s lab led to the development of a *Ca*^2+^ binding affinity prediction model using an amino acid sequence of EF-hand loop of CBPs. They have generated position-specific scoring metrics of EF-hand motif sequences to find the correlation between sequence and binding affinities.^19,23^ Franke *et al.* derived a simple linear model to estimate the correlation between experimental *Ca*^2+^-binding affinity with binding free energy calculation of crystal structures of CBPs using Fold-X and AutoDock vina methods. ^24^ Research conducted more than two decades ago by Boguta *et al.* developed simple rules to estimate the *Ca*^2+^ binding constant of the EF-hand domain based on the conformational properties of a given domain with the ideal reference pattern.^25^ There have also been studies that probed the mechanisms for altered binding affinities.^26–28^ Earlier studies carried out by our research group led to the development of a regression model for *Ca*^2+^ binding free energy calculation, which provides accurate predictions for calmodulin. ^29,30^ However till now, no generalized *Ca*^2+^ binding affinity prediction model tailored specifically for CBP variants has incorporated sequence, and structure along some specific properties that affect their interactions.

In our current study, we have developed an integrative modeling approach using the essential structure and sequence characteristics in conjunction with *Ca*^2+^ interaction properties of a CBP, cardiac Troponin-C, and its different variants. Our model has been built in the following manner. First, an accurate molecular dynamics (MD) based protocol has been used to calculate the *Ca*^2+^ binding free energy (BFE) of mutations using the MM-PBSA (Molecular Mechanics Poisson-Boltzmann Surface Area)^31^ method. The effect of polarization in the MD simulation is included by a charge scaling procedure. The protocol has been optimized using the experimental binding affinity data for the observed mutants of cardiac troponin-C (cTn-C). These mutants are primarily disease mutations (DM) associated with various heart disorders. In the next step, we have augmented our mutation dataset by generating additional plausible mutations (PM) from the perspective of sequence evolution of cTn-C protein. The evolutionary information-based mutation prediction method EVmutation has been used for this purpose developed by Hopf *et al.*^32,33^ Next, we calculated the BFE for those plausible mutations using the same protocol we had used earlier, and the *Ca*^2+^ BFE dataset of all the mutations was collected. Finally, using this dataset, a machine learning (ML) model to predict the *Ca*^2+^ binding affinity has been developed. Following the refinement and tailoring process, the most effective ML model was selected based on a rigorous evaluation of the performance metrics of various algorithms. Support Vector Regression (SVR)^34^ algorithm has been deployed to predict the binding free energy of mutants relative to that of the wild-type using nine features (comprising sequence, structure, and *Ca*^2+^ interaction properties extracted from MD simulation) of the mutation dataset. Our optimized prediction model with polynomial kernel gives a coefficient of determination (*R*^2^) of 0.77, Mean Squared Error (MSE) of 0.16 kcal/mol along with Pearson’s correlation coefficient (*R_p_*) of 0.92. The favorable results of these evaluation metrics highlight the effectiveness of the model.

The present study outlines an integrative computational approach utilizing a physics-based strategy, an evolutionary approach using a statistical probability method, and a support vector machine algorithm, utilizing sequence, structure, and *Ca*^2+^ interaction features, for the prediction of *Ca*^2+^ binding affinity towards cTn-C variants. As the methodology is general, it can be used for any other CBPs and it is expected to be an important integrative approach for predicting and designing CBPs with tailor-made *Ca*^2+^ binding affinity.

## 2. Methodology

This sections succinctly outline the methodology deployed in this study along with its theoretical foundations and protocols used for the calculations presented herein.

### 2.1 Molecular dynamics simulation to calculate the binding free energy for the mutations having experimental binding affinities

#### 2.1.1 Implementation of charge scaling for *Ca*^2+^ and its coordinating ions in classical Force Field

Among the different interactions in biological systems, polarization effects play an important role in the interactions of charged ions such as *Ca*^2+^ and salts.^35,36^ However, the majority of biomolecular simulations opt for non-polarizable force fields due to the complexities and computational constraints of explicitly including polarization. However in the presence of polyvalent ions, non-polarizable force fields often lead to a pronounced overestimation of ion pairing, potentially distorting simulation outcomes.^36,37^ Leontyev and Marshall came up with a model, where the electronic part of the dielectric constant of the medium is modeled by a simple charge scaling procedure. ^37,38^ This method, within its approximations, has been used for both isolated ions in water and protein in water. The amount of scaling depends on the electronic part of the dielectric constant, which is not possible to determine exactly for a protein system. Following previous works, we scaled the charges of *Ca*^2+^, salt ions (*K*^+^, *Cl^−^*), and *Ca*^2+^ coordinating and non-coordinating specific oxygen atoms by a scaling factor of 0.75^30,37,39^ (details are in supplementary Table S1 and S2). Our main aim is to establish a computational procedure that gives a good agreement with experimental binding free energy.

#### 2.1.2 Molecular Dynamic Simulation and Parameters

All simulations in this study were performed using the AMBER20 package,^40^ where the GPU-enabled cuda program has been used for the production run and the pmemd.MPI program to perform the steps up to system equilibration. To set up a molecular dynamics simulation of the cTn-C protein, we started with the structure of the wild-type (WT) protein of the calcium-saturated N-terminal regulatory domain of human cardiac troponin-C obtained from the Protein Data Bank (PDB ID:1AP4)^41^ which is shown in Figure 1. This figure represents the N-lobe of Cardiac troponin-C protein containing one non-functional loop I and *Ca*^2+^ bound EF-hand loop II. The first EF-hand of cTnC is unable to bind *Ca*^2+^ because of a single residue insertion (Val28) and two chelating residue substitutions (Asp29 → Leu and Asp31 → Ala). Calcium binding in the EF-hand II of the 12 amino acid residue loop is facilitated by the involvement of six negatively charged residues. These residues are specifically located at positions 1 (x), 3 (y), 5 (z), 7 (-y), 9 (-x), and 12 (-z). This figure also depicts five helices marked as N, A, B, C, and D. The initial structure served as a template for the construction of mutant proteins by changing specific residues in the wild-type protein structure. The protonation state of the residue was determined at a neutral pH using the pdb2pqr server (https://server.poissonboltzmann.org/pdb2pqr).^42^ Missing atoms and hydrogens in the mutant protein were added using the tleap module. The all-atom ff14SB force field^43^ was used to describe system parameters and the system was solvated in a TIP3P water model containing 0.15 M of KCl.^44^ The minimum distance between the protein and the cubic box edge was set to 10Å. Periodic boundary conditions were applied throughout all simulations. For the monovalent and divalent ions, we have used the Joung-Cheatham force field parameters and Li-Merz force field parameters respectively.^45,46^ Once the appropriate parameters set were established, we commenced by minimizing the system through 5000 steps using the steepest descent method, followed by an additional 5000 steps of the conjugate gradient method. During this phase, the protein was restrained with a force constant of 2 kcal/mol/Å^2^ on the heavy atoms of protein and *Ca*^2+^. Subsequently, we proceeded to relax the entire system for another 10,000 steps. van der Waals and short-range electrostatic interactions were truncated at a spherical cutoff distance of 10Å and long-range electrostatic interactions were accounted for by the particle mesh Ewald (PME) summation algorithm.^47^ SHAKE algorithm^48^ was used to constrain the bond distances involving hydrogen atoms with a time step of 2 fs for integrating the equation of motion. Following the minimization process, we gradually increased the temperature of the system to reach 300 K and kept it constant using a Langevin thermostat^49^ with a collision frequency of 2*ps^−^*^1^ and the Berendsen barostat^50^ with a pressure relaxation time of 1ps, was utilized for constant pressure. After 10 ns of heating and the subsequent 10 ns of density equilibration, the system underwent a 10 ns of isothermal-isobaric (NPT) equilibration. Next, we have run ten different trajectories of 10ns each for MM-PBSA calculations.^31^ Decorrelated configurations from simulation trajectories were saved using the block averaging method^51,52^ (details given in the supplementary information and represented in Figure S1). Cpptraj^53^ program was used to process and analyze the trajectory files. VMD visualization tool^54^ tcl script was used to calculate an average number of water molecules coordinating within 3^°^*A* of *Ca*^2+^ ion during simulation.

**Figure 1:**
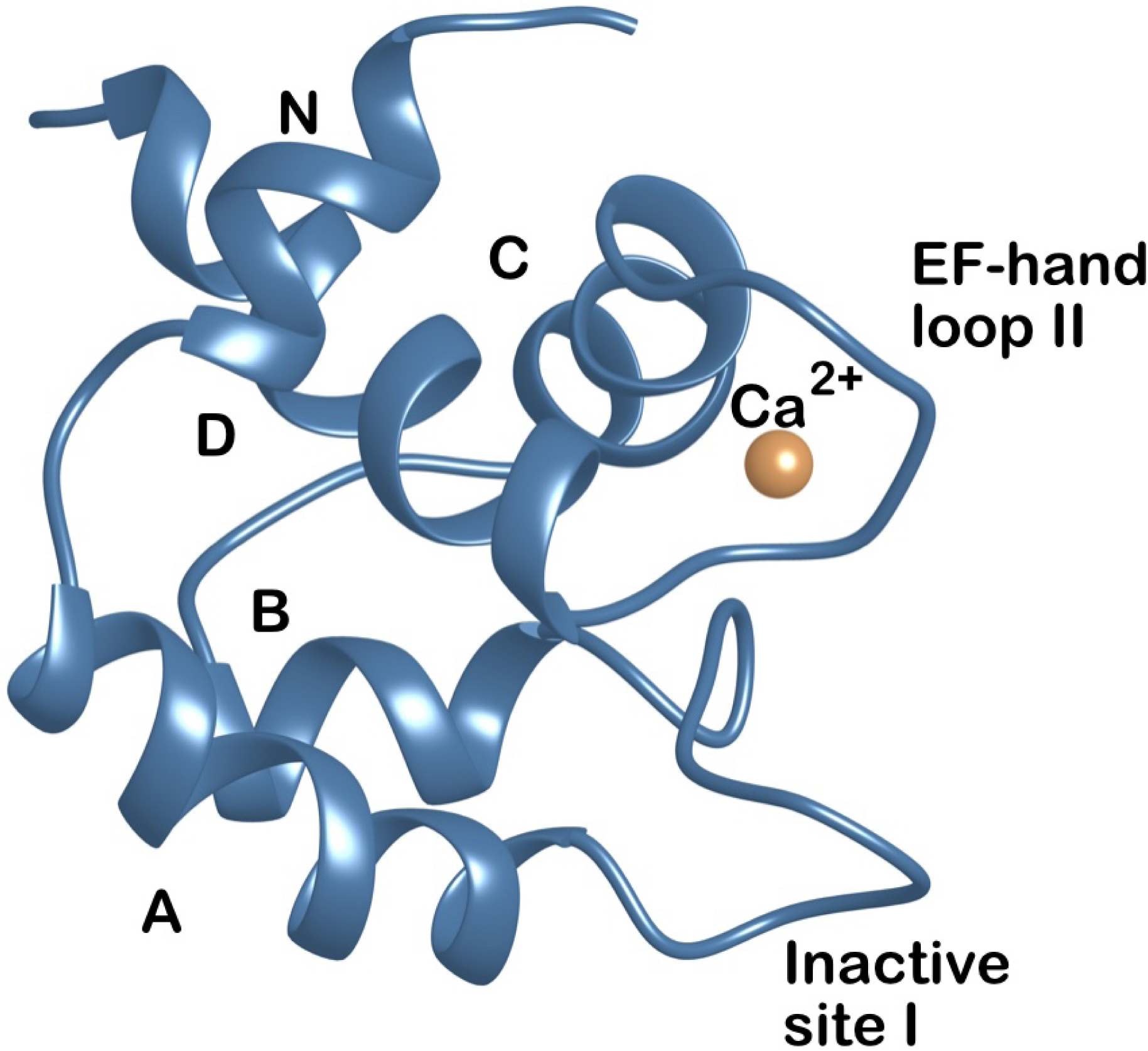
Cartoon representation of the N-lobe of cardiac troponin-C protein with *Ca*^2+^ ion present in the EF-hand loop II (PDB id: 1AP4). The figure also represents five helices depicted by N,A,B,C,D respectively with one inactive loop I.

#### 2.1.3 BFE Calculation using the MM-PBSA method

Total binding free energy (Δ*G_bind_*) of the protein-ion complex in the MM-PBSA method is approximated as:

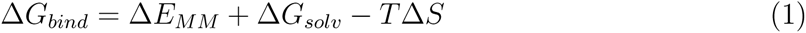

where Δ*E_MM_* represents a change in the gas-phase interaction and Δ*G_solv_* is the change in solvation-free energy upon ion binding. Δ*E_MM_* is calculated as the sum of electrostatic charges and van der Waals interactions upon ion binding. The decomposition of the energy term Δ*E_MM_* in MM-PBSA is given as:

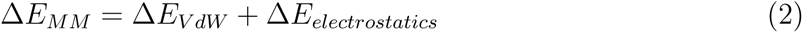

The solvation-free energy (Δ*G_solv_*) consists of two components related to ion binding as:

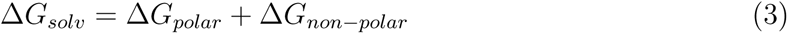

The calculation of Δ*G_polar_* using Poisson-Boltzmann equation involves estimating the electrostatic part of the solvation energy in the protein-ion complex upon ion binding. The decomposition of non-polar energy terms into cavitation and dispersion is given as:

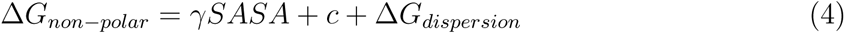

Here *γ* is the proportionality constant representing the change in free energy per unit molecular surface area and SASA is the Molecular surface area of the protein-ion complex. Here constant terms are c = −0.569 kcal/mol and *γ* is 0.0378 kcal/mol *A*^2^ used for our calculations.^55^ The entropy component (-TΔ*S*) is not considered in our calculations.

The calculations Δ*G_bind_* using the MMPBSA.py^56^ program were conducted with the ionic strength of 0.15 M and the dielectric constant of 78.35 for water. The dielectric constant of the protein was optimized to provide the most accurate agreement between our calculations with experimental observations. Relative BFE (ΔΔ*G_bind_*) of all the mutations have been calculated at various dielectric constants of protein ranging from 8 to 14 and we found that a dielectric constant of 12 gives the lowest RMSD between calculated and experiment ΔΔ*G_bind_* as shown in Figure 2.

**Figure 2:**
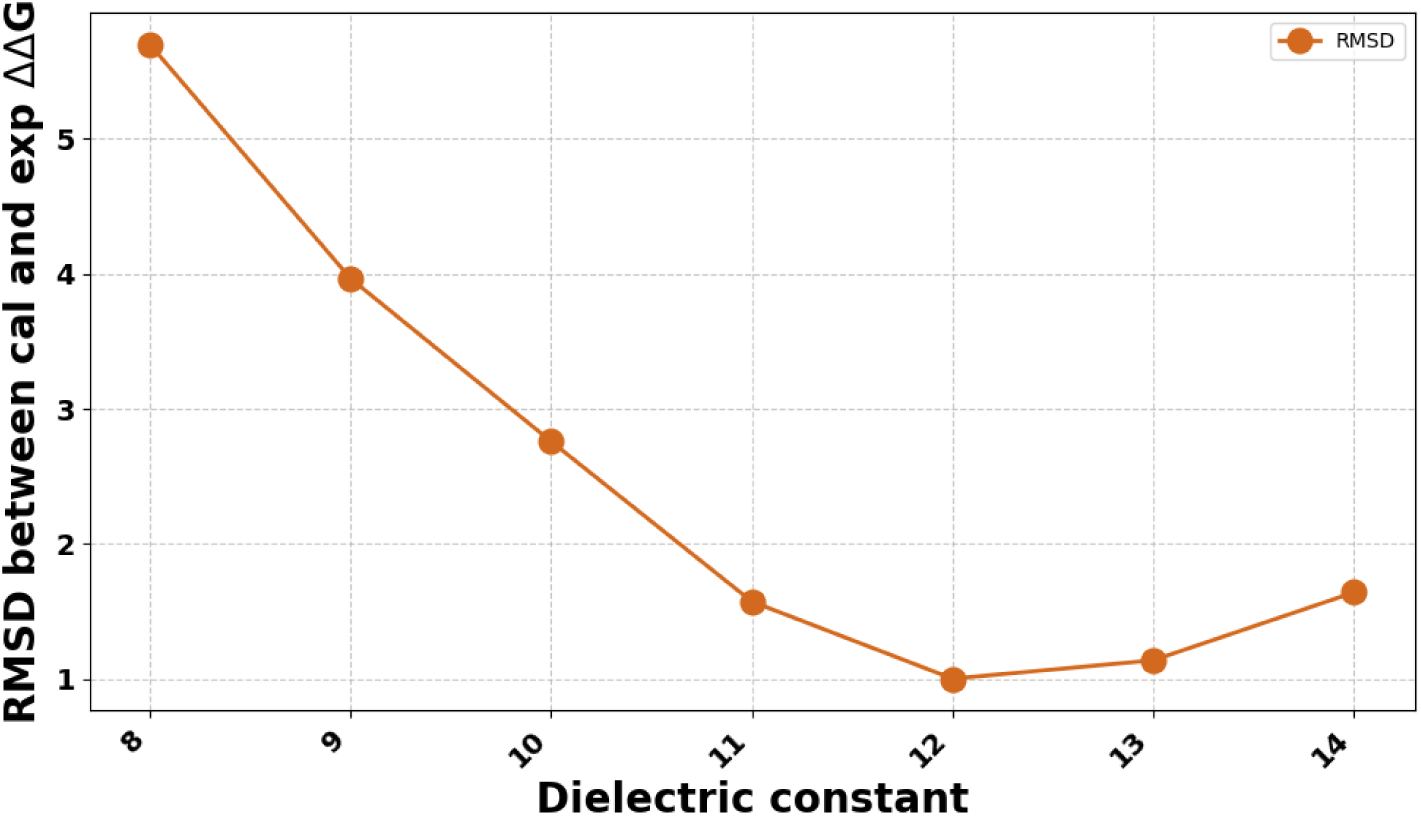
RMSD between calculated (at different internal dielectric constants) and experimental relative binding free energy, ΔΔ*G*. Relative binding free energy is the difference between mutants and the wild-type, which can be represented as ΔΔ*G* = Δ*G_mutant_* - Δ*G_W T_*. The internal dielectric constant of 12 gives the lowest RMSD.

### 2.2 Plausible mutations through an evolutionary approach using statistical probability method

A theoretical deep mutational scan along the sequence of cTn-C protein was done using a statistical probability method “EVmutation” developed by Hopf et al.^32^ This method includes both site-specific constraint and pairwise constraints between two amino acid positions i and j, represented in Figure 3.^57^

**Figure 3:**
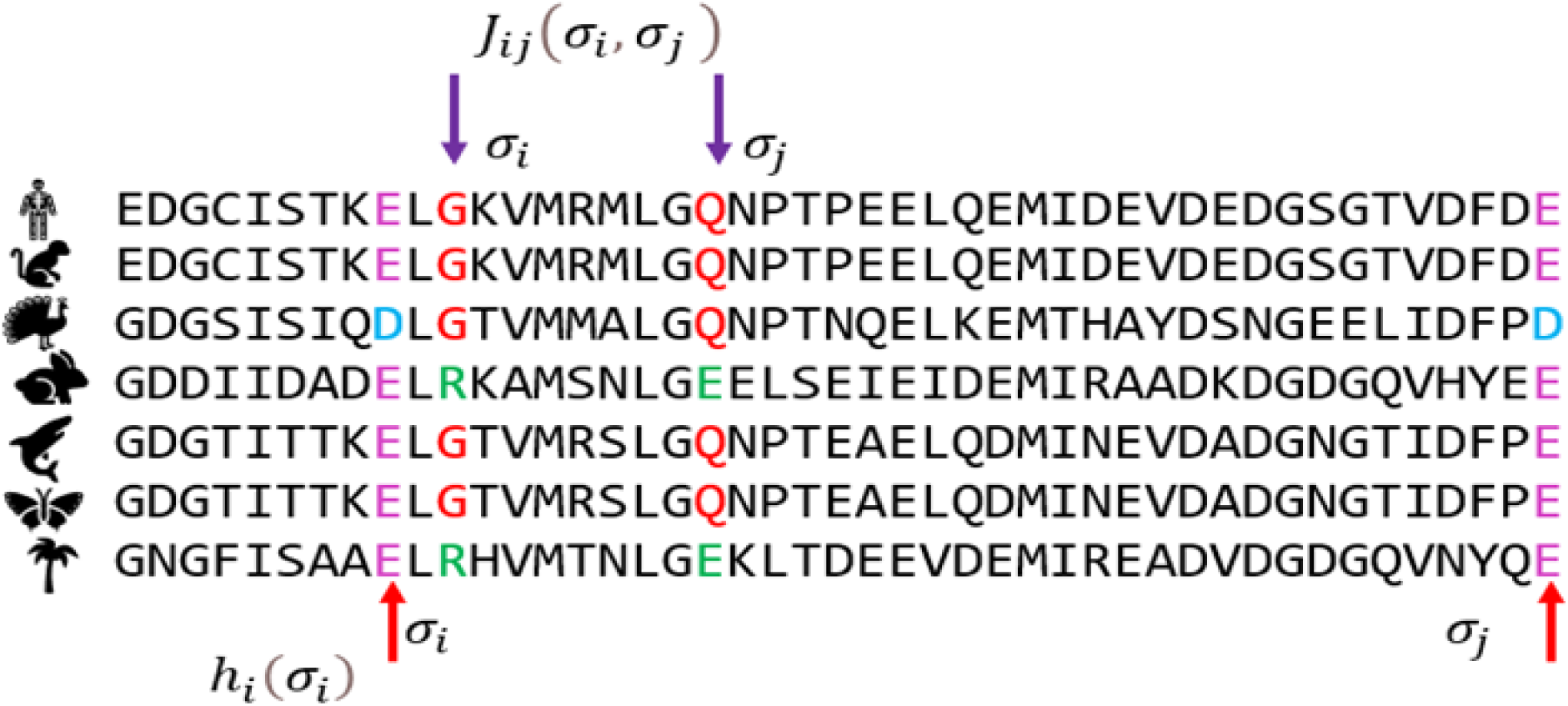
Schematic representation of multiple sequence alignment of a cardiac troponin-C protein sequence with site-specific constraints (*h_i_*) and coupling constraints (*J_ij_*) between residues i and j within sequence *σ*.

The total energy of specific sequence E(*σ*) is given by

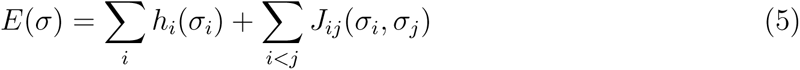

Here, E(*σ*) is an energy function used to capture different types of constraints in the sequence. *h_i_* represents site-specific constraints of amino acid at the position i to determine the likelihood of each amino acid residue within a sequence *σ* of length N. *J_ij_*(*σ_i_, σ_j_*) represents the coupling strength between residues i and j within a sequence *σ*. This coupling strength denotes the degree to which substituting an amino acid at the *i^th^* position influences the fitness of an amino acid at the *j^th^* position, and vice versa.

Using these parameters, the effect of each possible substitution has been quantified by evaluating the log-odds ratio of sequence probabilities (P) between the wild-type and mutant sequence as evolutionary statistical energy change (Δ*E_stat_*) as shown below

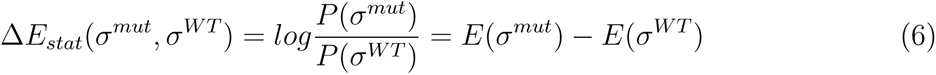

This EVmutation model incorporates the sequence context into the calculation of mutation effects.

#### 2.2.1 Selection of plausible mutations using EVmutation

In our study, we used the cardiac troponin-C sequence with Pfam id: PF13833 and Uniprot id: P63316, to access the quantitative effect of mutations. Multiple sequence alignment of the protein was done using the Jackhammer tool using the UniRef100 database and a length-normalized bit score threshold of 0.3 was applied.^32,33^ We have identified 76,931 homologous sequences of the target protein. To enhance the quality of the sequences, a post-processing step was performed to filter out positions with more than 30% gaps and sequences showing less than 50% alignment with the target sequence length and the complexity of the model applying *l*_2_ regularization. The EVmutation model has been performed and the results were analyzed, the evolutionary statistical energy change for each mutation (Δ*E_stat_*) was extracted from the protein mutational landscape. For the selection of plausible mutations (PM), the value of Δ*E_stat_* was analyzed. Δ*E_stat_* value greater than zero indicates that mutant sequences are more probable and potentially beneficial. In contrast, values below zero suggest that mutant sequences are less probable and possibly deleterious. Based on this criterion, we chose sixty-one beneficial mutations. We also extracted the evolutionary conservation score for each residue along the sequence length. The conservation score is a measure to identify how different parts of a protein are important. Conserved positions remain the same across different species of the protein and are usually critical for the protein to be functional, while nonconserved parts are more likely to change. Essentially, a mutation in the residue having a high conservation score is more likely to be damaging.

### 2.3 Machine Learning Model for predicting the binding affinity

The machine learning (ML) model utilized for the prediction of *Ca*^2+^ relative binding free energy(ΔΔ*G_pred_*), where ΔΔ*G_pred_* = Δ*G_mut_* - Δ*G_W T_* denoted as:

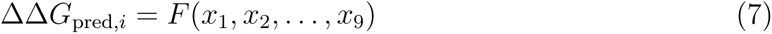

where F is the ML model adopting a nonlinear function, and *x*_1_*, x*_2_*, …x*_9_ are the nine input features forming a feature vector.

#### 2.3.1 Benchmarking Input Feature Selection

The gist of machine learning lies in generating distinctive input features. Our model includes simple sequence, structure, and *Ca*^2+^ interaction properties and is composed of nine features that are systematically classified based on their intrinsic characteristics.

##### Sequence-based features

We have included four properties: a) conservation score of each residue along the protein sequence, b) evolutionary statistical energy difference upon mutation (Δ*E_stat_*), c) position of the mutation in the protein sequence, and d) change in the charge of a protein due to a mutation. The conservation score represents the importance of conserved amino acid residue during sequence evolution, which may be critical for the stability of protein structure and function.

##### Structure-based features

We have included four properties: a) type of amino acid substitution, b) secondary structure of the residue at the mutation site, c) distance of *Ca*^2+^ from the center of mass of the mutant residue, and d) change in SASA (solvent-accessible surface area) of mutant residue from WT residue (*SASA_W T_* − *SASA_mutant_*). The physicochemical differences between wild-type and mutant residues were quantified by categorizing amino acids into four groups: hydrophobic (Gly, Ala, Val, Leu, Ile, Met, Cys, Pro), aromatic (Phe, Trp, Tyr, His), polar (Ser, Thr, Asn, Gln), and charged (Asp, Glu, Lys, Arg). Then each group was encoded by assigning 1,2,3,4, respectively. The type of amino acid substitution describes the properties of amino acids that change upon mutation, for example: charged to polar is encoded as 1 (4-3), and hydrophobic to polar as −2 (1-3), etc. The secondary structure of the residue at the mutation site indicates whether the mutation took place in helix, sheet, coil, or turn. SASA was calculated to quantify the exposure of amino acid to the solvent and thus determine its accessibility to the molecular surface. The SASA values of each amino acid are presented in Table S3.^58^

##### Ca^2+^ interaction properties

It includes the average number of water molecules coordinating within 3^°^*A* radius of *Ca*^2+^, during the simulation. The number of water molecules present in the vicinity of *Ca*^2+^ changes the coordination shell of *Ca*^2+^ binding loop and thus affects the *Ca*^2+^ binding affinity.

#### 2.3.2 Machine learning models

##### Prediction model

Several ML models were used to predict *Ca*^2+^ relative binding free energy of cTn-C protein variants. Support vector regression, decision tree, and random forest algorithms were used to build the prediction model and evaluate its performance. This comparative analysis aims to identify the best prediction algorithm customized for our mutation dataset. Out of these three algorithms, Support Vector Regression (SVR) exhibits superior performance and thus most effective model for our dataset. SVR conducts regression tasks by seeking an optimized hyperplane that minimizes the sum of distances between data points and the hyperplane. In our study, we used SVR using the scikit-learn library,^59^ where input features were standardized using the StandardScaler to ensure uniform scaling. The mutation data set was randomly split into training and test sets using a split ratio 80%-20%, respectively. Hyperparameter tuning was conducted using GridSearchCV, exploring various combinations of hyperparameters, such as the regularization parameter (*C*), margin of tolerance (*ɛ*), and type of kernel used. Following tuning of optimized hyperparameters, the model was trained on the training set and the performance of the model was carried out on the test set using evaluation metrics including Mean Squared Error (MSE), coefficient of determination (*R*^2^), and Pearson’s correlation coefficient (*R_p_*). Next, we have performed 5-fold cross-validation^60^ to estimate our model’s performance more reliably. This approach helps identify any potential overfitting by estimating performance over multiple training and test splits. We have utilized Stratified K-Fold Cross-Validation to evaluate the performance and generalization capability of the SVR model. This method was carefully chosen due to the unequal distribution of two major classes in our data set that contain positive and negative values of target variable. Stratified K-Fold ensures that each fold in the cross-validation process retains the same proportion of each class as in the original dataset, minimizing the potential for bias due to class imbalance.

##### Assessing the contribution of each feature

In the succeeding steps, we carried out an analysis to identify the contribution of each input feature in the prediction model. For this purpose, we have applied three different approaches to discern the features that contribute most significantly to the model. First, we calculated the correlation coefficients of each feature with ΔΔ*G_bind_* using linear, polynomial, and radial basis function kernels. Then we have used the Permutation feature importance method^61^ which assesses the impact of each feature on model performance by measuring the change in accuracy when the values of that feature are randomly shuffled. To investigate the role of each feature, we have used the SHAP (SHapley Additive exPlanations) method^62^ which provides a unified measure by assigning a contribution score to each feature for the prediction of the model, based on SHapley values. For this, we have calculated Shapley values by considering all possible ways in which the features can be combined, giving a measure of how much each feature contributes. The use of these approaches for the evaluation and analysis of each feature enhances the interpretability and transparency of the model.

## 3. Results and Discussions

### 3.1 Binding Free Energy Calculations

The charge scaling protocol with the MM-PBSA method has been used to calculate the *Ca*^2+^ BFE of cTn-C protein mutations. We have collected *Ca*^2+^ binding affinity experimental data of twenty-four mutations available in the literature as represented in Table 1. The mutation sites in the protein structure are represented in Figure S2. Binding free energy calculations have been done for this dataset and the mean value of Δ*G_bind_* and its different components, namely Δ*E_vdW_*,Δ*E_elec_*, Δ*E_PB_*,Δ*E_cavitation_*, and Δ*E_dispersion_* are summarized in Table S4. Relative binding free energy calculations are performed to accurately assess how mutations influence *Ca*^2+^ binding affinity relative to the wild-type protein. We have used multiple short trajectories to calculated the mean of the BFE and mean of standard error of the mean, which would indicate the deviation in the BFE. This approach has been previously validated by our group and as well as by different authors to calculate the average BFE from independent multiple but short trajectories.^29,63^ Relative binding free energy is represented as ΔΔ*G_bind_*, where ΔΔ*G_bind_* = Δ*G_mutant_* - Δ*G_W T_* has been calculated for mutations and depicted in Figure 4a, to observe the alignment between calculations and experiments. This figure depicts that all (except F20Q mutant) the relative binding free energy calculations are within 0.87 kcal/mol to the experimental relative BFE. Our observation reveals the following patterns: a) negatively charged residue substitution to any other amino acids reduces the binding affinity of *Ca*^2+^. This effect is more prominent when the mutation happens in the loop region while it shows less noticeable effect when the mutation is located outside the loop. For example, E40A, D67A, D73A, and D75Y mutants present in the loop show a significant reduction in *Ca*^2+^ binding affinity (positive ΔΔ*G_bind_*) whereas D87A and D88A mutations located outside the loop do not show much difference. This finding indicates that beyond the type of amino acid substitution (for instance, polar to non-polar mutation), the location of the mutation within the three-dimensional structure is also crucial, b) hydrophobic to glutamine amino acid substitution was seen in F20Q, A23Q, L29Q, V44Q, L48Q, I61Q, V79Q, and M81Q and these mutations show a consistent increase in *Ca*^2+^ binding affinity (negative ΔΔ*G_bind_*) irrespective of the position of the mutation except L57Q. This finding showcases that non-polar to polar specifically glutamine substitution increases binding affinity, c) Some mutations including Y5H, S37G, E59D, I61Q, D87A, and D88A show very low ΔΔ*G_bind_* values, reflecting that they are almost neutral in terms of their effect on *Ca*^2+^ binding, d) E59D and I61Q mutations appear as an opposite trend between calculated and experiment ΔΔ*G_bind_* although the difference is not significant in E59D.

**Figure 4:**
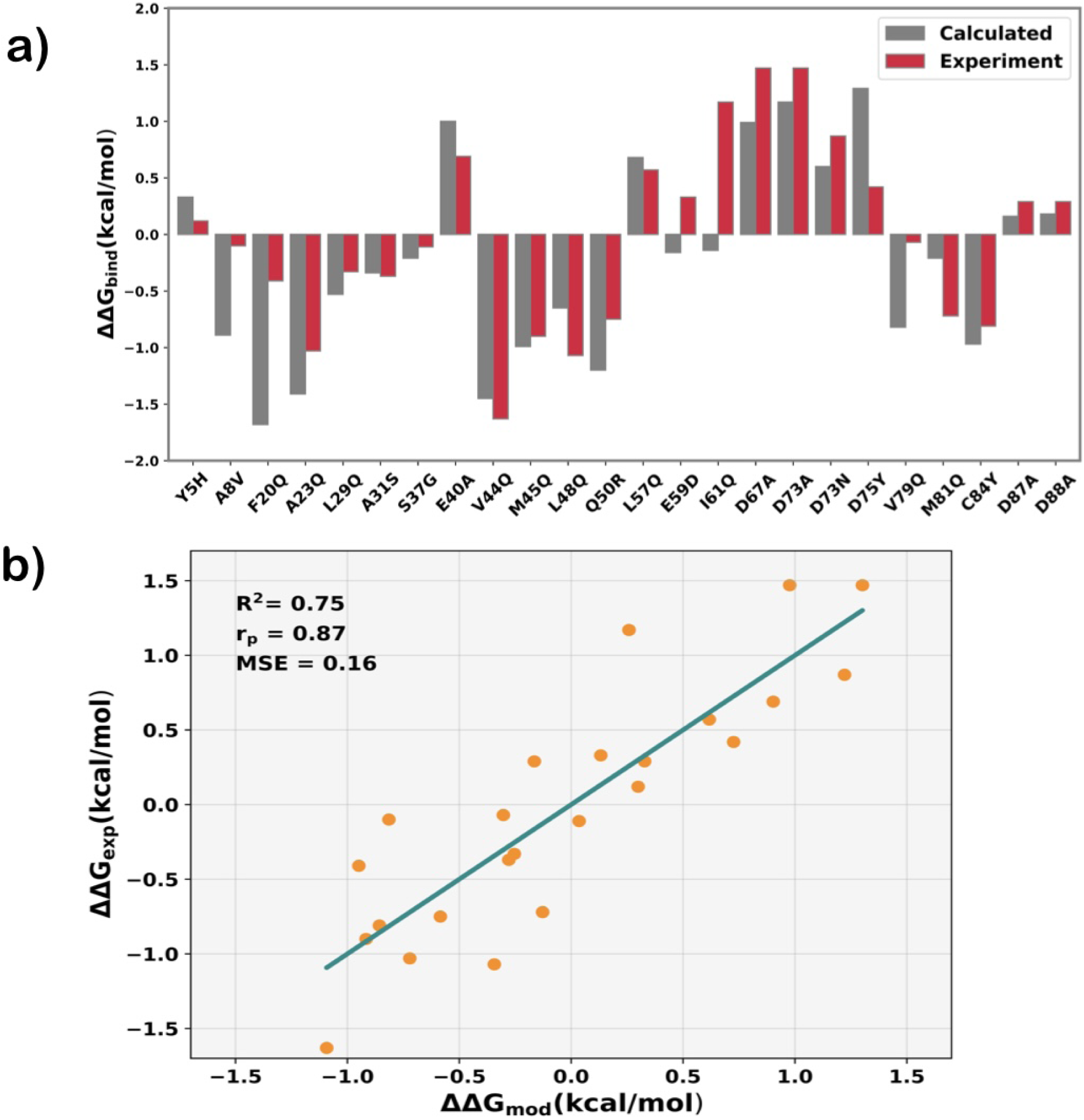
a) The barplot for the comparison between the calculated relative binding free energy (grey) and their corresponding experiment values (red) (in kcal/mol) for twenty-four mutations. On the x-axis, the first and the last letters represent the wild-type and mutant amino acid respectively, and the number represents the position of that amino acid in the protein sequence. b) The scatter plot between modeled ΔΔ*G_mod_* and experimental ΔΔ*G_exp_* relative binding free energy for mutants. The solid line represents the best linear fit for the dataset. The coefficient of determination (*R*^2^), Mean Squared Error (MSE), and Pearson’s correlation coefficient (*R_p_*) are also shown.

**Table 1:** List of cardiac troponin-C wild-type (WT) protein and diseased mutants with their experimental binding affinity towards calcium. The names of mutants are represented in which the first and last letters denote a single-letter code representation of the amino acid of wild-type and mutant respectively, and the numbers between these two letters indicate the mutated position. Δ*G_exp_* value with standard error(in kcal/mol) is presented in this table.

| System | $\Delta G_{exp}(kcal/mol)$ |
| --- | --- |
| WT | -6.56 (0.03)<sup>8</sup> |
| Y5H | -6.44 (0.02)<sup>6</sup> |
| A8V | -6.66 (0.01)<sup>8</sup> |
| F20Q | -6.97 (0.06)<sup>17</sup> |
| A23Q | -7.59 (0.13)<sup>9</sup> |
| L29Q | -6.89 (0.03)<sup>8</sup> |
| A31S | -6.93 (0.06)<sup>8</sup> |
| S37G | -6.67 (0.03)<sup>9</sup> |
| E40A | -5.87 (0.13)<sup>9</sup> |
| V44Q | -8.19 (0.05)<sup>17</sup> |
| M45Q | -7.46 (0.02)<sup>17</sup> |
| L48Q | -7.63 (0.01)<sup>8</sup> |
| Q50R | -7.31 (0.06)<sup>8</sup> |
| L57Q | -5.99 (0.02)<sup>10</sup> |
| E59D | -6.23 (0.01)<sup>13</sup> |
| I61Q | -5.39 (0.02)<sup>10</sup> |
| D67A | -5.09 (0.06)<sup>18</sup> |
| D73A | -5.09 (0.06)<sup>18</sup> |
| D73N | -5.69 (0.08)<sup>5</sup> |
| D75Y | -6.14 (0.06)<sup>13</sup> |
| V79Q | -6.63 (0.05)<sup>9</sup> |
| M81Q | -7.28 (0.03)<sup>17</sup> |
| C84Y | -7.37 (0.12)<sup>8</sup> |
| D87A | -6.27 (0.01)<sup>12</sup> |
| D88A | -6.27 (0.01)<sup>12</sup> |

#### Regression model for accurate prediction of ΔΔG_exp_

Moving forward, our next step involves quantifying the accuracy of our calculated ΔΔ*G_cal_* with experimental ΔΔ*G_exp_* as shown in Table 2. Figure 4b depicts a linear regression model using the Ordinary Least Square module of sklearn in Python based on BFE components as descriptors for the prediction of the BFE (details are given in supplementary information). The regression model gives Pearson’s correlation coefficient (*R_p_*) of 0.87 and *R*^2^ of 0.75 with MSE of 0.16 kcal/mol reflecting the robust performance of the model. To ensure the reliability of our model, we have performed leave-one-out cross-validation. The leave-one-out cross-validation validation gives the Pearson’s correlation coefficient (*R_p_*) of 0.78, *R*^2^ of 0.60, and MSE of 0.26 kcal/mol, which validates the reliability and predictive power of our model.

**Table 2:** This table lists the calculated relative binding free energy depicted as ΔΔ*G_cal_* and experimental relative binding free energy shown as ΔΔ*G_exp_* of cardiac troponin-C mutations (in kcal/mol).

| System | $\Delta\Delta G_{cal}$ | $\Delta\Delta G_{exp}$ |
| --- | --- | --- |
| Y5H | 0.33 | 0.12 |
| A8V | -0.89 | -0.1 |
| F20Q | -1.68 | -0.41 |
| A23Q | -1.41 | -1.03 |
| L29Q | -0.53 | -0.33 |
| A31S | -0.34 | -0.37 |
| S37G | -0.21 | -0.11 |
| E40A | 1.00 | 0.69 |
| V44Q | -1.45 | -1.63 |
| M45Q | -0.99 | -0.9 |
| L48Q | -0.65 | -1.07 |
| Q50R | -1.2 | -0.75 |
| L57Q | 0.68 | 0.57 |
| E59D | -0.16 | 0.33 |
| I61Q | -0.14 | 1.17 |
| D67A | 0.99 | 1.47 |
| D73A | 1.17 | 1.47 |
| D73N | 0.6 | 0.87 |
| D75Y | 1.29 | 0.42 |
| V79Q | -0.82 | -0.07 |
| M81Q | -0.21 | -0.72 |
| C84Y | -0.97 | -0.81 |
| D87A | 0.16 | 0.29 |
| D88A | 0.18 | 0.29 |

#### Residue-wise binding free energy decomposition

Residue-wise BFE decomposition of loop II residues has been done to check the contribution of coordinating and non-coordinating residue to BFE. Figure 5 shows the results for WT and a few mutants. From this observation, we can ascertain that negatively charged amino acid residues present at the 1*^st^*,3*^rd^*,9*^th^*, and 12*^th^* positions show a significantly high contribution to BFE in the WT system. Therefore, to understand the effect of mutations on the contribution of these residues we compare them with WT. Concerning F20Q mutation, aspartic acid at the 3*^rd^* position exhibits a significantly higher contribution compared to the wild-type (WT) and for V79Q and M81Q mutation, aspartic acid at the 3*^rd^*, 9*^th^*, and 12*^th^* position shows a much higher contribution than WT, hence leading to a higher ΔΔ*G_bind_*. Conversely, for the D75Y mutation, the contribution of all the negative charge residues is markedly reduced resulting in a substantial decrease in the ΔΔ*G_bind_*.

**Figure 5:**
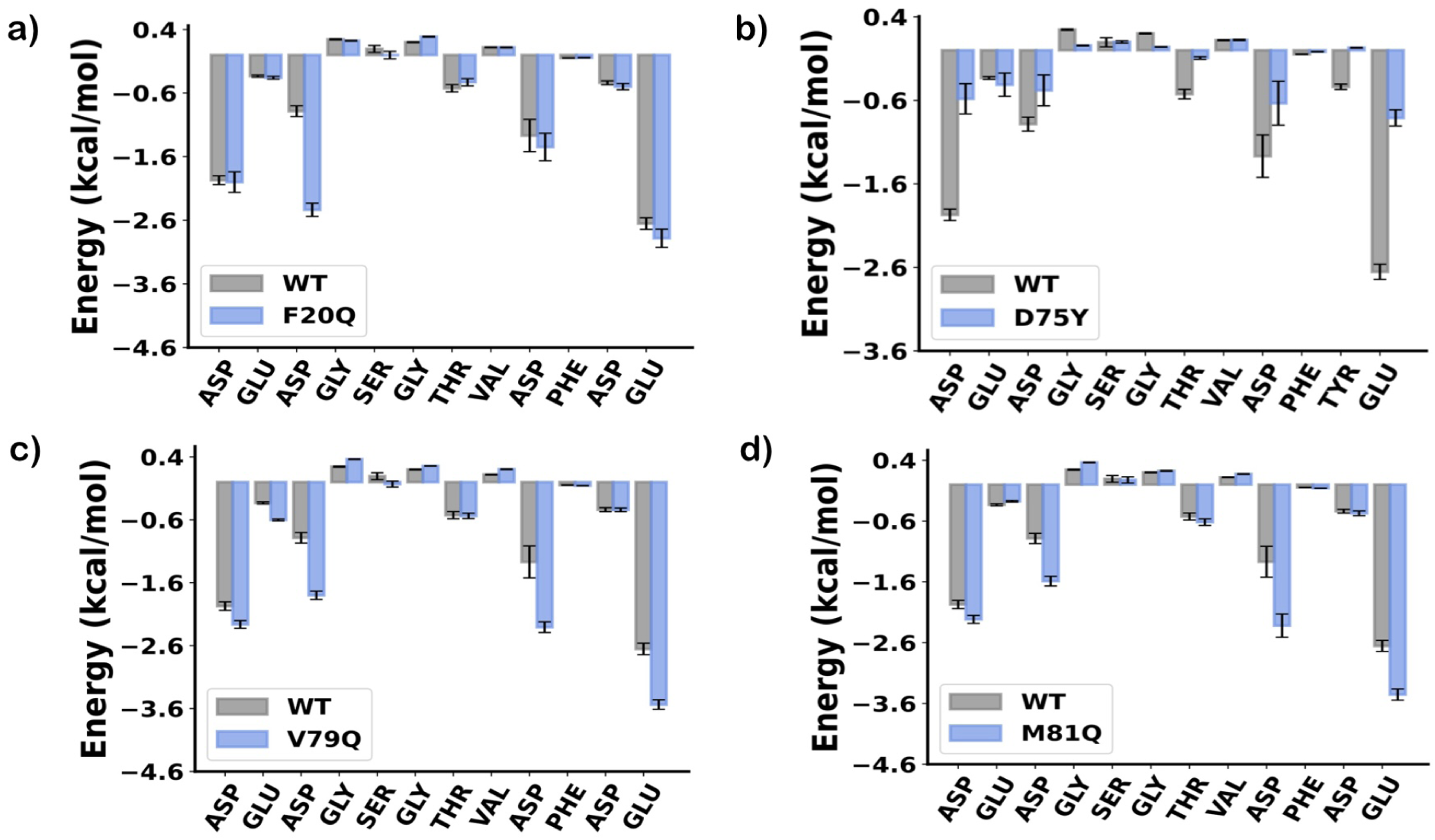
The barplot for residue-wise binding free energy decomposition of EF-hand loop II residues (12 residues). This figure illustrates the contribution of individual residue to the binding free energy in the wild-type (grey) protein and four key mutants (blue), specifically F20Q, D75Y, V79Q, and M81Q. The error bars represent the standard error of the mean.

#### Role of water in coordination sphere

The presence of water molecules in coordination with *Ca*^2+^ has a significant impact on the BFE. In the WT protein, one water molecule stays coordinated with the *Ca*^2+^ ion throughout the simulation. Results for a few mutations are shown in Figure 6. From the figure, it can be seen that M45Q and V79Q mutations (which have higher binding affinity than the WT) have one and zero coordinating water molecule, respectively. On the other hand, the D75Y mutation, which causes a decrease in binding affinity, shows the presence of three coordinating water molecules with *Ca*^2+^ leading to disruption of the coordination sphere.

**Figure 6:**
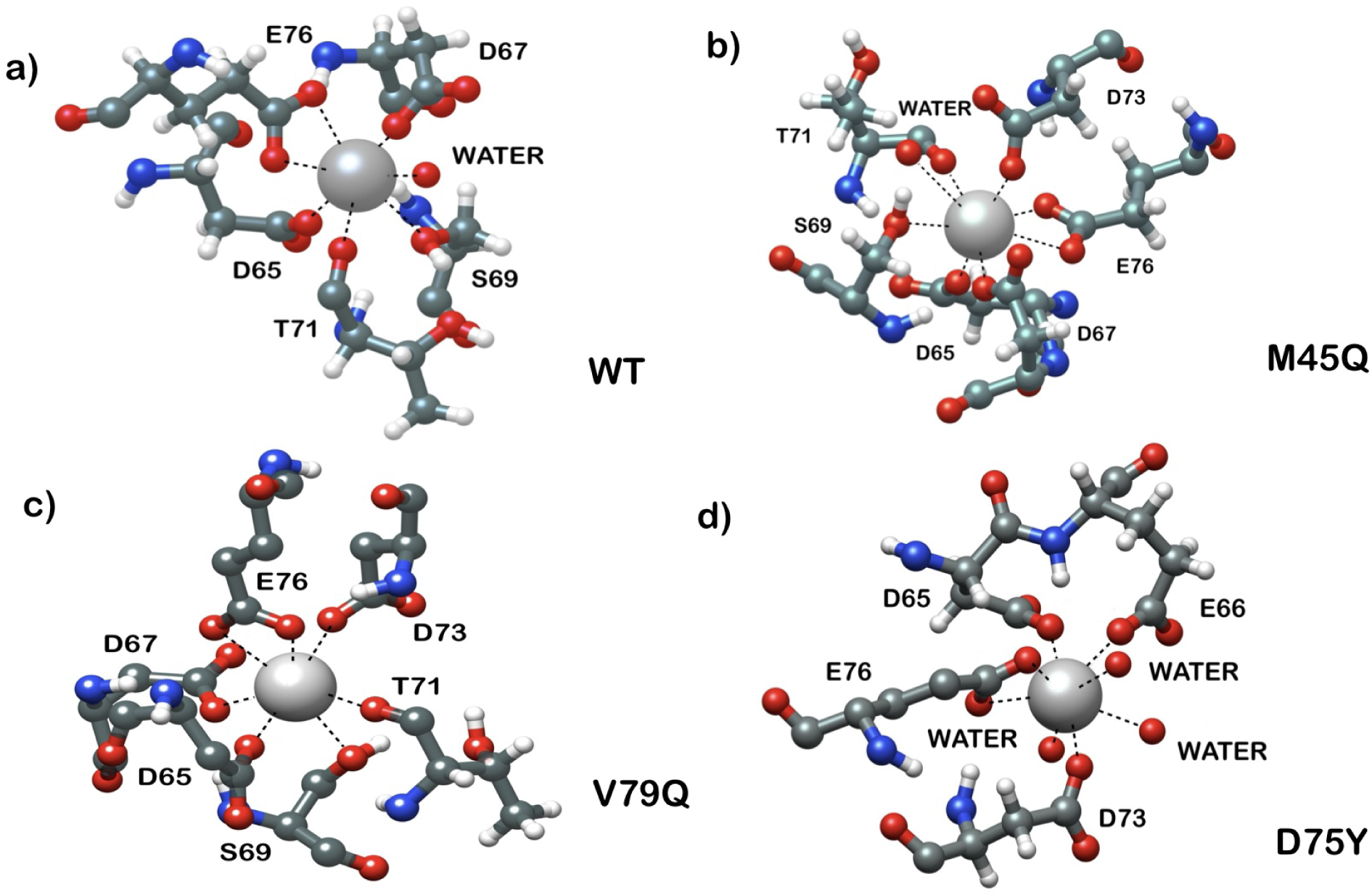
This figure represents *Ca*^2+^ coordinating oxygen atoms of EF-hand loop II residues and water in the (a)wild-type cardiac troponin-C protein and its mutants, (b) M45Q, (c) V79Q, and (d) D75Y. In the WT system and M45Q mutant, one water is coordinating with *Ca*^2+^ and three water molecules are present in D75Y mutant and no water is present in V79Q mutant.

### 3.2 Generation of plausible mutations using evolutionary information

We augmented the mutation dataset by generating additional plausible mutations using the EVmutation method described in the method section. The cardiac troponin C sequence has been used as a target sequence to evaluate the impact of each mutation in terms of evolutionary statistical energy difference (shown in equation 6). Figure 7a represents the mutational landscape of evolutionary statistical energy difference on each mutation in the protein sequence (from residues 18-81). The evolutionary fitness of each mutation showcased through the lens of Δ*E_stat_*. Each square in the mutational landscape denotes Δ*E_stat_* value in terms of color gradient representing beneficial and deleterious mutations, as indicated by the Δ*E_stat_* scale. Hence, we selected mutations having Δ*E_stat_* ranging from 0 to +4 and the highest Δ*E_stat_* value was predominantly observed in mutations namely G30D, G30N, L29D, L29E, A22E, M47S, P54D, P54E and followed by other mutations. In contrast, Δ*E_stat_* ranging from 0 to −12 represented deleterious mutations such as mutations concentrated in residues D65X, D67X, L41X, F24X, F77X, and many more (here X represents any other 19 amino acids except in WT). To gain insights into the discrete evolutionary behaviors of beneficial and deleterious mutations, we generated individual plots for each group to separate the mutations with positive Δ*E_stat_* (beneficial) from those with negative Δ*E_stat_* shown in Figure 7b. This plot highlights beneficial and deleterious mutations shown in teal and orange color dots providing two distinct visualizations. Subsequently, we have selected sixty-one beneficial mutations depicted in the subplot for inclusion in the PM dataset.

**Figure 7:**
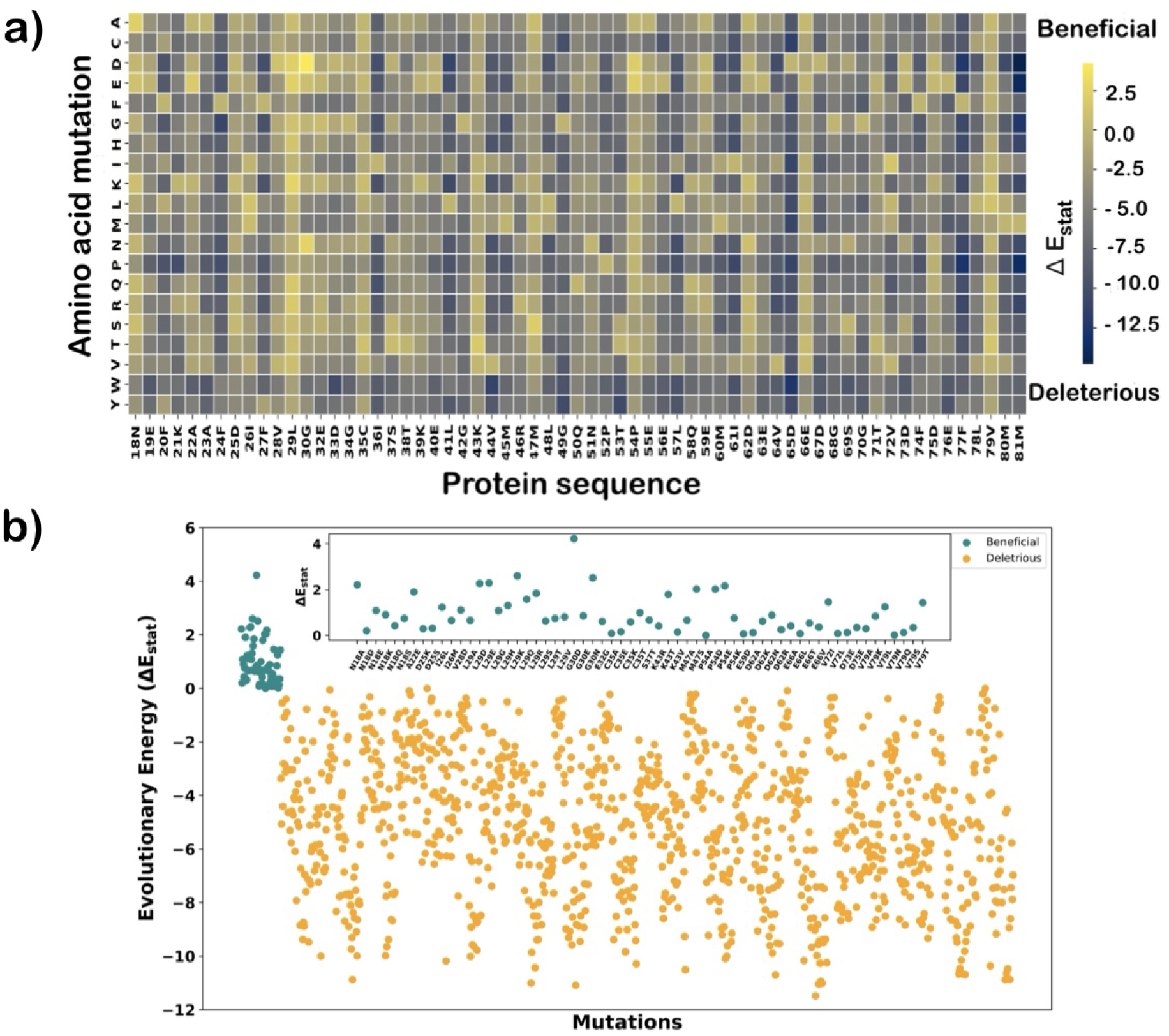
(a) The heatmap showing the evolutionary energy change upon each mutation represented by Δ*E_stat_*. On the x-axis, the number represents the residue position and the letter is a single code representation of the amino acid present at this position. The color gradient from yellow to blue reflects the Δ*E_stat_* values of each mutation in the landscape in terms of beneficial and deleterious mutation respectively. (b) The scatter plot of the evolutionary energy change upon each mutation, Δ*E_stat_* for all mutants. The beneficial (positive values) and deleterious (negative values) are shown in teal and orange colors respectively. In the subplot, mutants having positive Δ*E_stat_* are represented.

Proceeding with the conservation score of each residue in the protein sequence aiding in the identification of critical regions that are essential for protein function and *Ca*^2+^ binding activity. Focusing on the conservation pattern reveals that residues having a high conservation score are mostly present in the *Ca*^2+^ binding loop region while residues present outside the binding loop show a low conservation score, illustrated in Figure S4. Loop residues from 29 to 40 (loop I) and 65-76 (loop II) are mostly conserved except for a few.

BFE calculations of PM have been done using the MM-PBSA protocol described before. To develop a machine learning-based BFE prediction model for cTN-C mutations, we have calculated the relative BFE (ΔΔ*G_cal_*) of PM and values are presented in Table S5. Our dataset for the ML consists of both DM and PM sets. Three mutations including L29Q, E59D, and V79Q are common in DM and PM, and six DM including Y5H, A8V, A31S, C84Y, D87A, and D88A do not have evolutionary information, hence these nine mutations are not considered in building the ML model.

### 3.3 Machine Learning Model to Predict BFE of Mutants

We have used an advance machine learning model, Support Vector Regression, for the prediction of *Ca*^2+^ binding affinity of cTn-C protein mutations.

As detailed in the Methods section, we have used nine different features for the prediction of BFE in SVR model represented in Table 3. For the model training process, the mean value of ΔΔ*G_cal_* of the mutation dataset has been used as the benchmark. Employing hyperparameter tuning on the mutation dataset, it is found that a margin of tolerance (*ɛ*) value of 0.5 and regularization parameter (C) of 1, with a polynomial kernel gives the best evaluation statistical parameters. Our SVR model with polynomial kernel gives Pearson’s correlation coefficient of 0.92 and *R*^2^ of 0.77, along with MSE of 0.16 kcal/mol, as depicted in Figure 8. Also, we have checked the SVR model on linear and rbf kernels with the same values of *ɛ* and C, and statistical parameters obtained using various kernels are shown in Figure S5. To eliminate the potential overfitting and make over model more reliable, we have performed Stratified 5-fold cross-validation. The cross-validation gives *R*^2^ of 0.75, *R_p_* of 0.88, and MSE of 0.27 kcal/mol. The results of the Stratified K-Fold Cross-Validation demonstrate that the model’s performance is consistent across all five folds. These findings highlight the robustness of the model.

**Figure 8:**
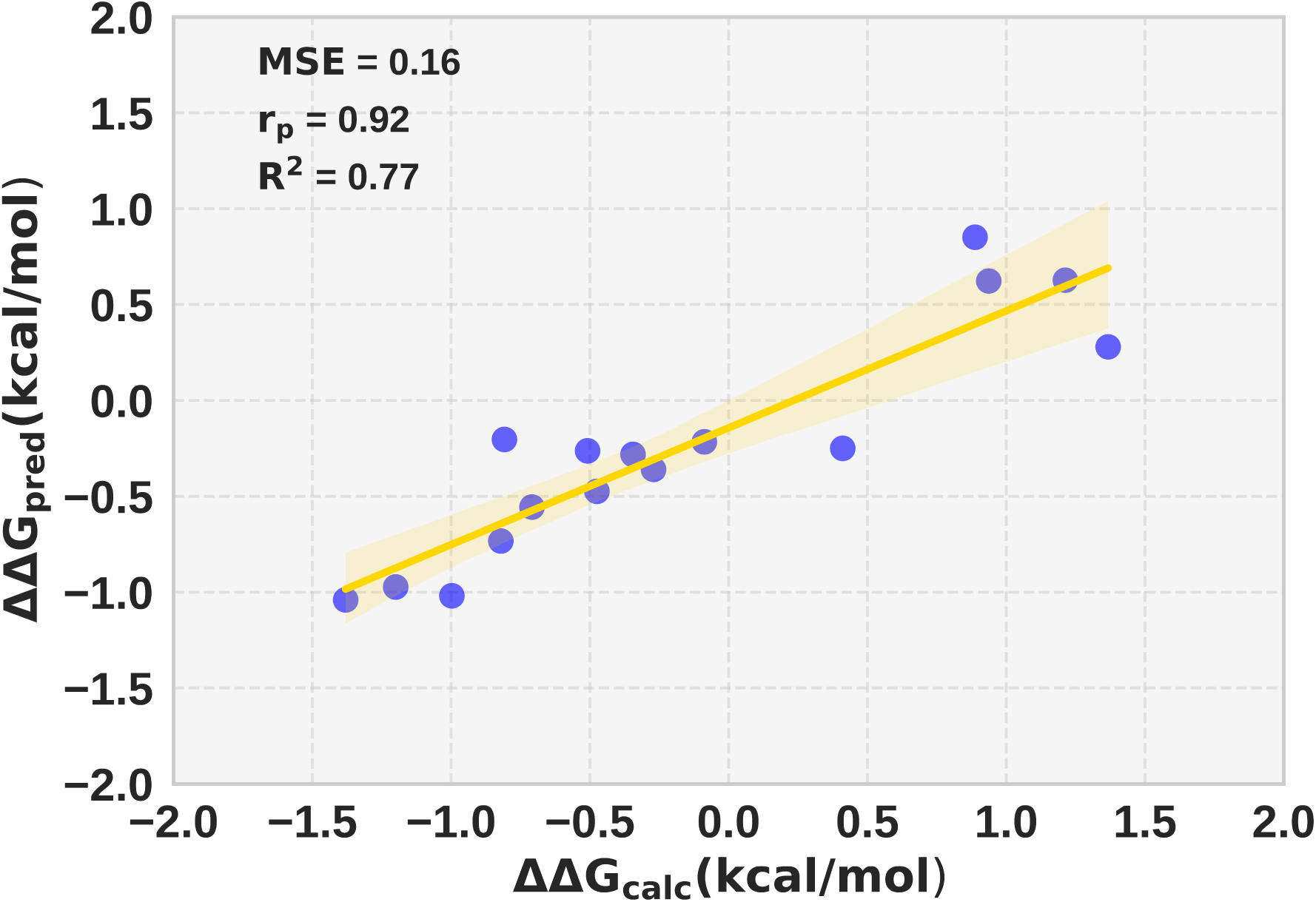
The scatter plot between the calculated relative binding free energy (ΔΔ*G_calc_*) and predicted relative binding free energy ΔΔ*G_pred_* on the test set (from the whole dataset) using a polynomial support vector regression model. The solid line represents the best fit. The coefficient of determination (*R*^2^), Mean Squared Error (MSE), and Pearson’s correlation coefficient (*R_p_*) are also shown.

**Table 3:** This table represents different feature types including sequence, structure, and *Ca*^2+^ interaction properties used in the machine learning model. Sequence and structure feature type covers four features each while *Ca*^2+^ interaction features type includes only one feature.

| Feature type | Features |
| --- | --- |
| Sequence | Conservation score of residue<br>Evolutionary statistical energy<br>Charge of the system<br>Position of mutation in protein sequence |
| Structure | Secondary structure of mutant site<br>Distance of mutation from $Ca^{2+}$<br>change in solvent accessible surface area<br>Type of amino acid substitution |
| $Ca^{2+}$ interactions | Number of water within $3\text{\AA}$ of $Ca^{2+}$ |

#### 3.3.1 Assessing the contribution of each feature

Machine learning algorithms are capable of mapping complex data and deriving interpretable relationships between observable and target variables. In this study, we have examined the importance of features by using correlation coefficients of each feature, permutation feature importance, and the SHAP analysis method. First, we calculated the correlation coefficients of each feature with ΔΔ*G_bind_* using three different kernels in the SVR (see Figure S6 in SI). The figure showcases that the number of water feature is the most positively correlated with ΔΔ*G_bind_*, while the distance with *Ca*^2+^ and the amino acid substitution features are the most negatively correlated withΔΔ*G_bind_*. The second method, permutation feature importance, permutes each feature and evaluates the impact on the accuracy of the model providing a clear understanding of how different features affect outcomes. Results obtained from this method in terms of the ranking of each feature contribution in the model are presented in Figure 9. This analysis confirms that the three features, number of water, amino acid substitution, and distance with *Ca*^2+^ collectively contribute the most around 80% (40%+25%+15%) to the prediction of the model. Further, we have employed the SHAP method to interpret and compare the impact of individual features on the overall performance of the model. The SHAP summary plot is shown in Figure 10, in which each dot represents a single prediction for an individual feature. The spread of SHAP values from negative to positive indicates the strength and consistency of each feature. For instance, the number of water feature shows the widest spread from negative(−0.3) to positive (0.7) SHAP value. This shows the highest impact while the conservation score feature (at the bottom) shows the least impact on the predictability of the model. In this analysis also, the features the number of water and amino acid substitution are the two most important features confirming the first two methods. However, in SHAP analysis charge change feature is the third most contributing feature. From the outcomes of these methods, we found the top three prominent features, number of water, amino acid substitution, and distance with *Ca*^2+^. These results encouraged us to investigate how effectively each descriptor class (sequence, structure, and *Ca*^2+^ interactions) contributes to the predictability of the SVR model. For this, we have used different combinations of descriptor classes for the prediction, and their statistical metrics and parameters are depicted in Table 4. The above finding emphasizes that the structure-based descriptor is the most contributing class that gives the highest *R*^2^ of 0.57, *R_p_* of 0.79, and lowest MSE of 0.31 kcal/mol. This finding can be used as a valuable reference for protein engineering research in the future, providing crucial insights for developing more effective models. Overall, we have developed a model that is capable of predicting the binding free energy of a given cTn-C mutation based on some physical descriptors, with high accuracy. In the scope of future studies, our model can be used to design mutations in CBPs to achieve desirable *Ca*^2+^ binding affinity.

**Figure 9:**
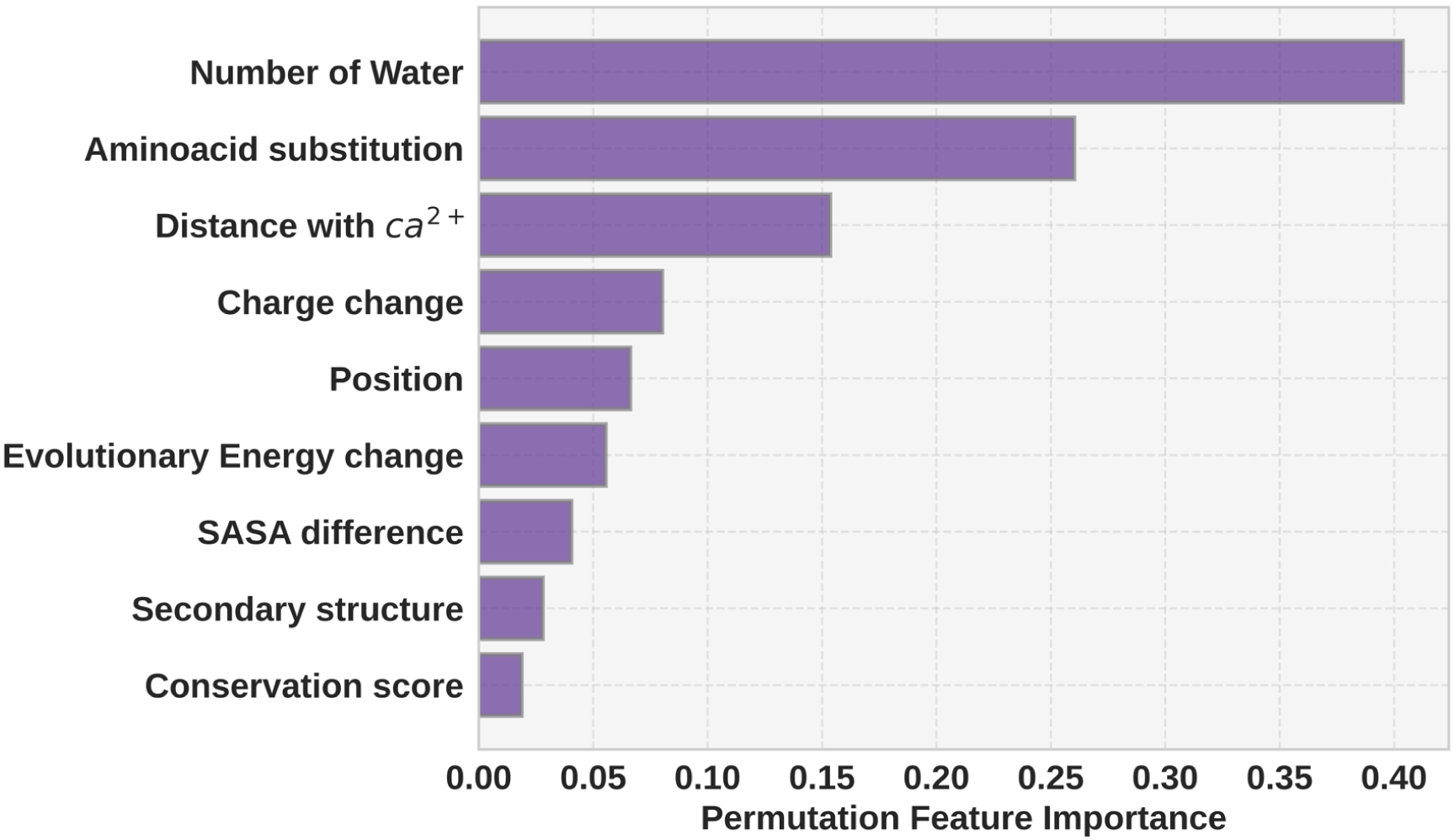
The permutation feature importance for the support vector regression model. The features are ranked according to their contribution to the model performance. The most important feature that appears at the top is the number of water, and the least important at the bottom is the conservation score.

**Figure 10:**
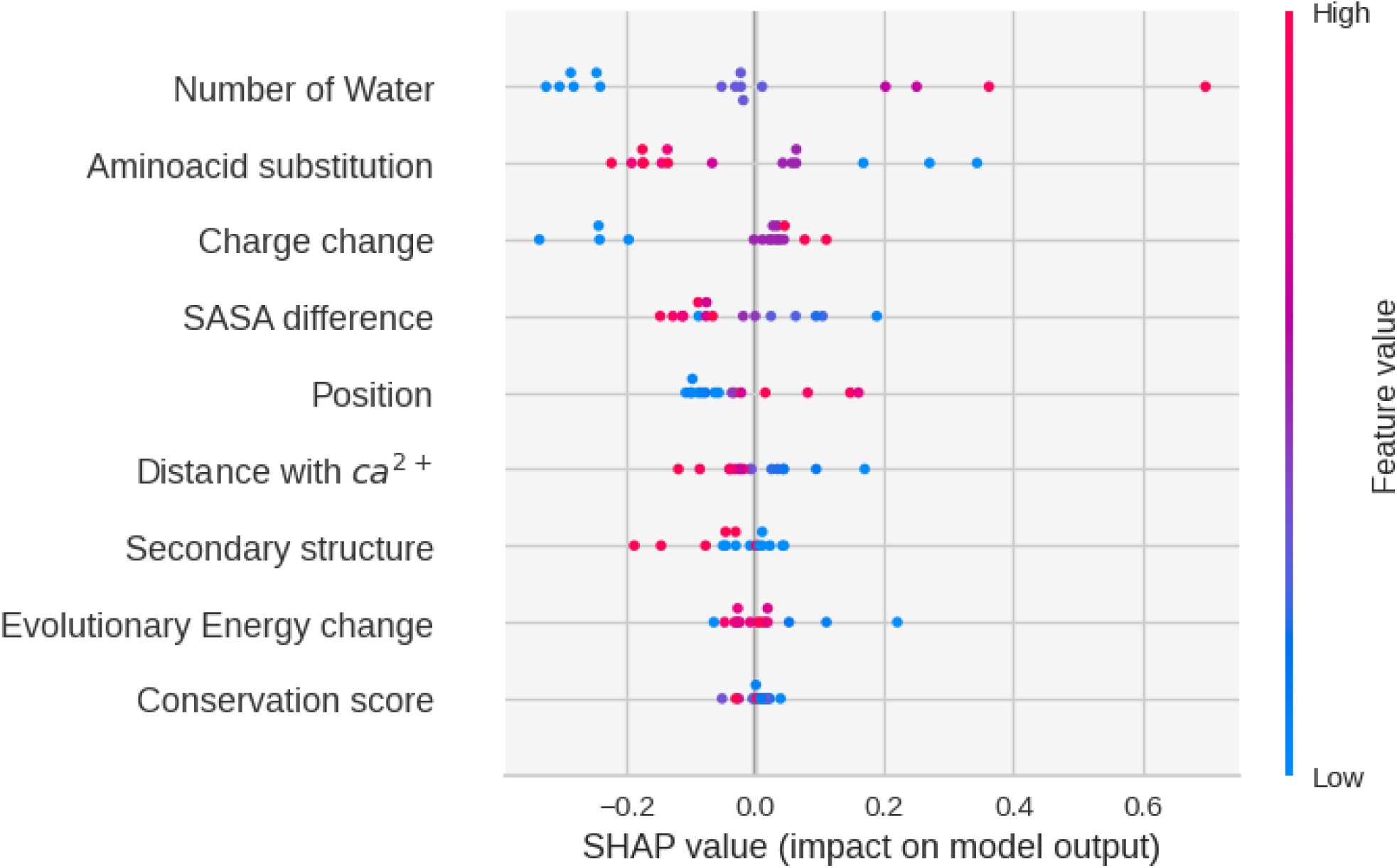
The beeswarm plot of the SHAP (SHapley Additive exPlanations) values of the features. Each dot represents the individual prediction and is colored according to the feature value. The trend of the color of the dots from left to light in blue to red indicates feature has a positive impact on SHAP values (model output) and vice-versa. The features at the top show the highest impact on the prediction of the model.

**Table 4:** Table shows a combination of different types of features with their evaluation metrics shown as coefficient of determination (*R*^2^), Mean Squared Error (MSE),Mean Absolute Error (MAE), Root Mean Square Error (RMSE), pearson’s correlation coefficient (*R_p_*) and best kernel type used in support vector regression model.

| Descriptors combination | $R^2$ | MSE | MAE | RMSE | pearson’s | kernel |
| --- | --- | --- | --- | --- | --- | --- |
| Sequence+Structure+$Ca^{2+}$ interaction | 0.77 | 0.16 | 0.30 | 0.41 | 0.92 | poly |
| Sequence+Structure | 0.51 | 0.35 | 0.51 | 0.59 | 0.80 | rbf |
| Sequence+$Ca^{2+}$ interaction | 0.66 | 0.25 | 0.44 | 0.50 | 0.81 | rbf |
| Structure+$Ca^{2+}$ interaction | 0.72 | 0.20 | 0.36 | 0.45 | 0.85 | rbf |
| Sequence | 0.34 | 0.47 | 0.57 | 0.69 | 0.59 | rbf |
| Structure | 0.57 | 0.31 | 0.44 | 0.56 | 0.79 | linear |
| $Ca^{2+}$ interaction | 0.47 | 0.38 | 0.54 | 0.62 | 0.69 | linear |

## 4. Conclusions

Predicting the binding affinities of *Ca*^2+^ binding to different calcium-binding proteins is an important yet non-trivial problem. The alteration of binding affinity upon mutation can have a profound effect on the function of CBP and can cause abrupt changes in CBP-dependent signaling pathways. Moreover, designing CBPs with tailor-made properties can be a valuable tool in synthetic biology. Our newly developed integrated modeling using molecular dynamics simulation, prediction of plausible mutations, and machine learning prediction for an important CBP, Troponin-C is an important step in predicting binding affinity between *Ca*^2+^ and a large number of mutants for the protein. Our model uses a combination of physics based and machine learning model and examines the evolution of Troponin-C for predicting plausible important mutants. In general, our model can be an important stepping stone for modeling the binding affinity in the mutational landscape of any CBP.

## Acknowledgement

This work is supported by a Department of Biotechnology (DBT) grant (BT/PR45865/BID/7/1016/2023) awarded to P.B. This work was also partially supported by grants from the DBT (BT/PR/40251/BITS/137/11/2021) awarded to the Centre for Computational Biology and Bioinformatics, Jawaharlal Nehru University and the MATRICS grant from SERB (MTR/2021/000365) awarded to P.B. The authors would like to acknowledge the help from Prof. S. Gourinath, Drs. Rakesh Mishra and Ajeet Kumar Yadav, and Mr. Abdul Basit. Pooja acknowledges CSIR (File No:09/0263(13722)/2022-EMR-I) for a senior research fellowship.

## Supporting Information Available

Detail of the parameters, additional plots, and data are given in Supporting Information.

