## Supplementary Information for "Calcium Binding Affinity in the Mutational Landscape of Troponin-C: Free Energy Calculation, Coevolution Modeling and Machine Learning"

### Estimating Statistical Errors

Simulation averages are calculated from a limited number of random samples, which can introduce imprecision in the estimation of averages and other statistical properties such as binding free energy, etc.[1] To provide estimates of the statistical uncertainties associated with the simulation averages and to allow the determination of the statistical significance value,[2] we have used the block averaging method to estimate statistical errors and assess the convergence of results in MD simulations. We have plotted the statistical inefficiency as a function of the block size in figure S1. The figure shows a plateau value at S=25. This suggests that in our simulation, roughly one configuration out of every 25 stored, contributes entirely novel information to the computed average.

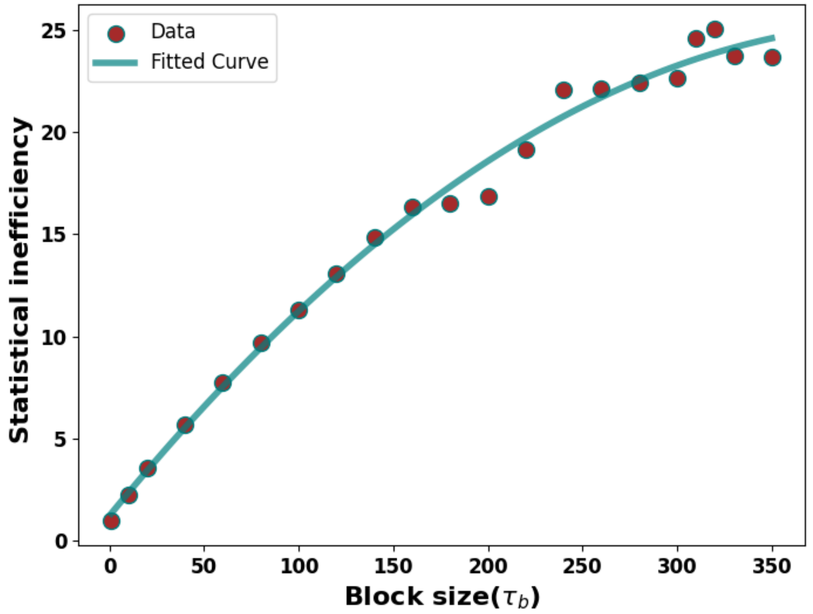


Figure S1: **Graphical plot illustrates the relationship between statistical inefficiency (S) against block size ($\tau_b$).**

#### Developed a Regression Model using Relative Binding free energy for diseased mutations

We expressed Relative Binding Free Energy (BFE) to quantify the difference between mutants and wild type $\Delta G_{bind}$ calculated as shown in equation:

$$\Delta\Delta G_{bind} = \Delta G_{mutant} - \Delta G_{WT} \tag{1}$$

Equation 8 represents a linear regression model used to estimate the change in binding free energy ($\Delta\Delta G_{\mathrm{mod}}$) between a mutant and a wild type (WT) protein based on various free energy components calculated in MM-PBSA, as a descriptor for our regression model.

$$\Delta\Delta G_{\mathrm{mod}} = b_0 + b_1\Delta\Delta E_{\mathrm{vdw}} + b_2\Delta\Delta E_{\mathrm{el}} + b_3\Delta\Delta G_{\mathrm{PB}} + b_4\Delta\Delta G_{\mathrm{cav}} + b_5\Delta\Delta G_{\mathrm{disp}} \tag{2}$$

This is shown in Figure 4b of the main text.

#### Figures and Tables

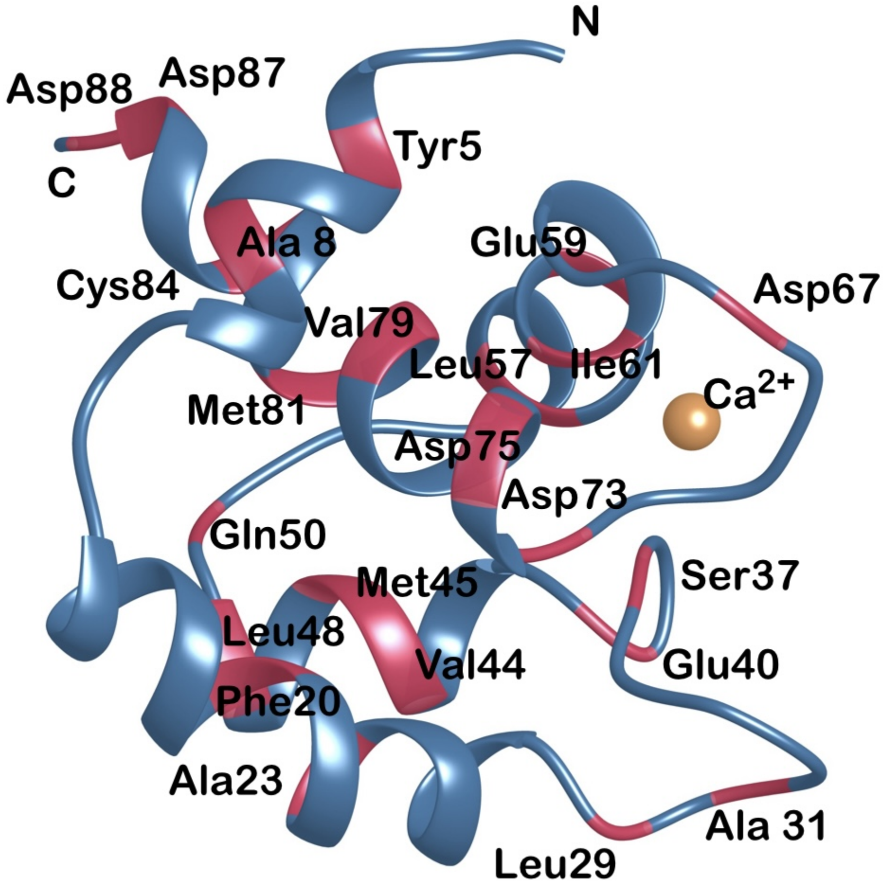


Figure S2: **This figure illustrates the diseased mutations site shown in red color on the N-lobe of cardiac troponin-C protein. N and C represent the N-terminal and C-terminal of the protein respectively. Mutants names are represented by a three-letter code of amino acid followed by the position of that amino acid in the protein sequence.**

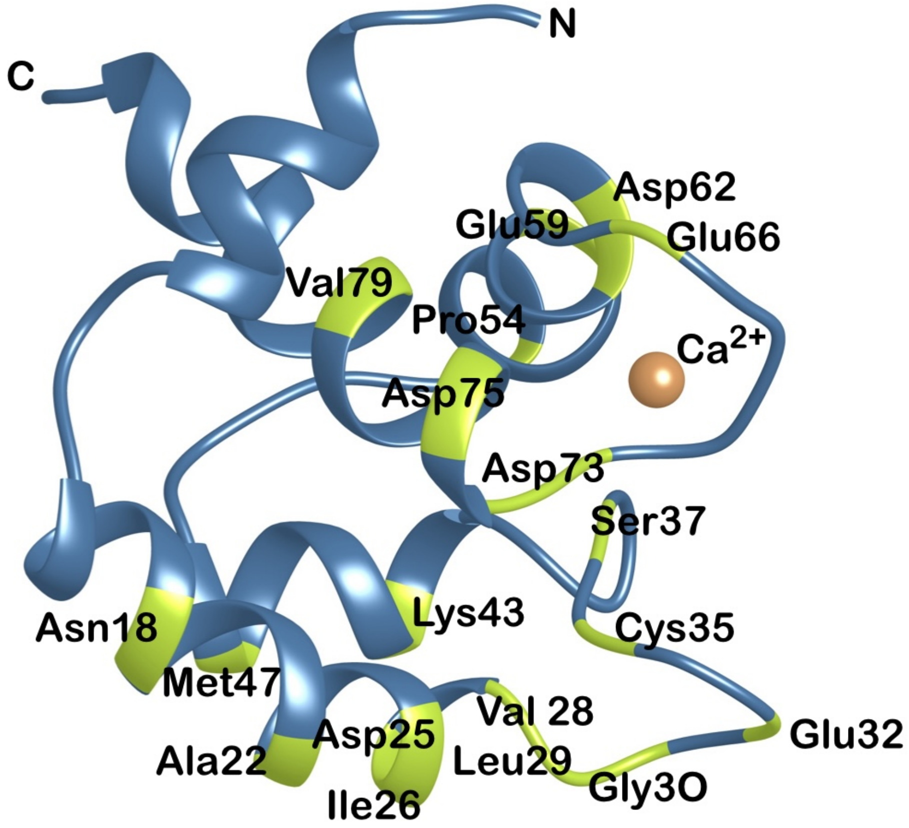


Figure S3: **This figure illustrates the site of the plausible mutations shown in lime green color on the N-lobe of cardiac troponin-C protein. N and C represent N-terminal and C-terminal of the protein respectively. Mutants names are represented by three-letter code of amino acid followed by the position of that amino acid in the protein sequence.**

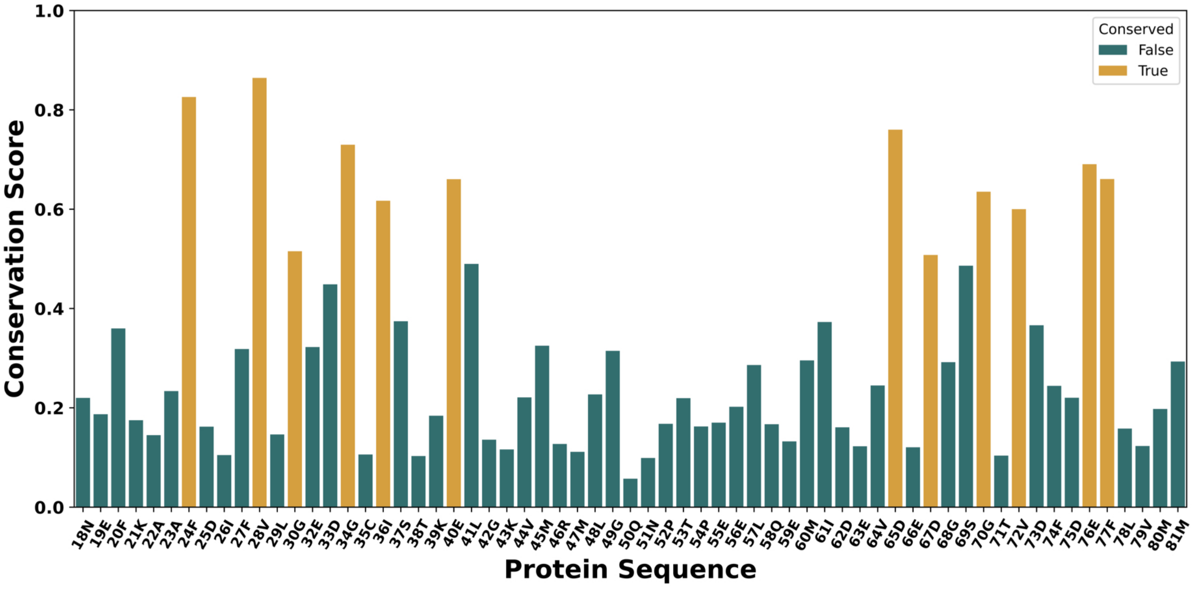


Figure S4: **Figure represents the conservation score of each residue along the protein sequence. The y-axis represents the conservation score from 0 to 1, in which 0 represents the least conserved and 1 represents the highest conserved residue. Residues with conservation scores greater than 0.5, are depicted by yellow bars, while green bars represent residues with lower conservation scores.**

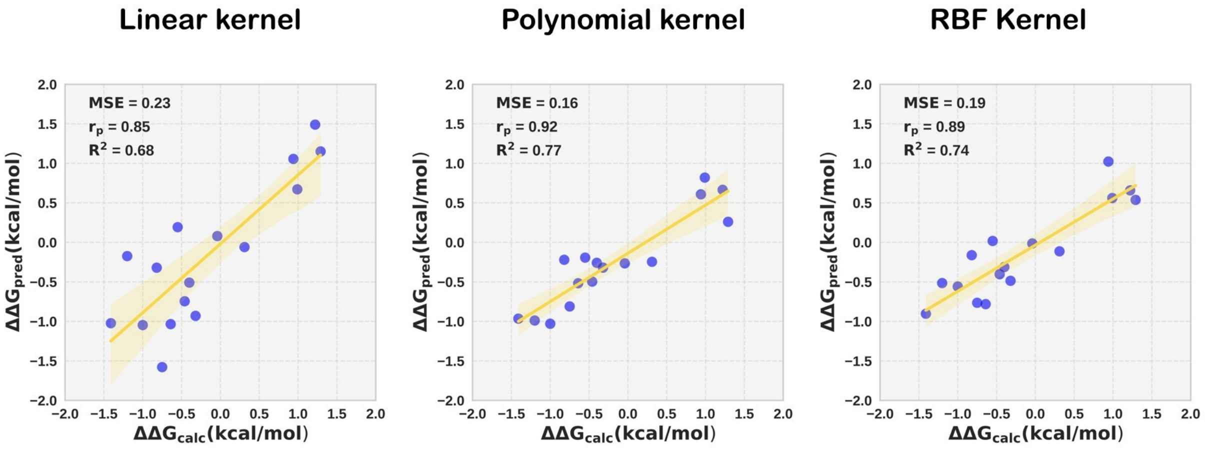


Figure S5: **This figure depicts a comparative plot using linear, polynomial, and rbf kernel (left to right) in the support vector regression model. The x-axis represents the calculated relative binding free energy ($\Delta\Delta G_{calc}$) while the y-axis represents predicted relative binding free energy $\Delta\Delta G_{pred}$. Evaluation metrics coefficient of determination ($R^2$), Mean Squared Error (MSE), and Pearson's correlation coefficient ($R_p$) are shown in the upper left corner of the individual plot.**

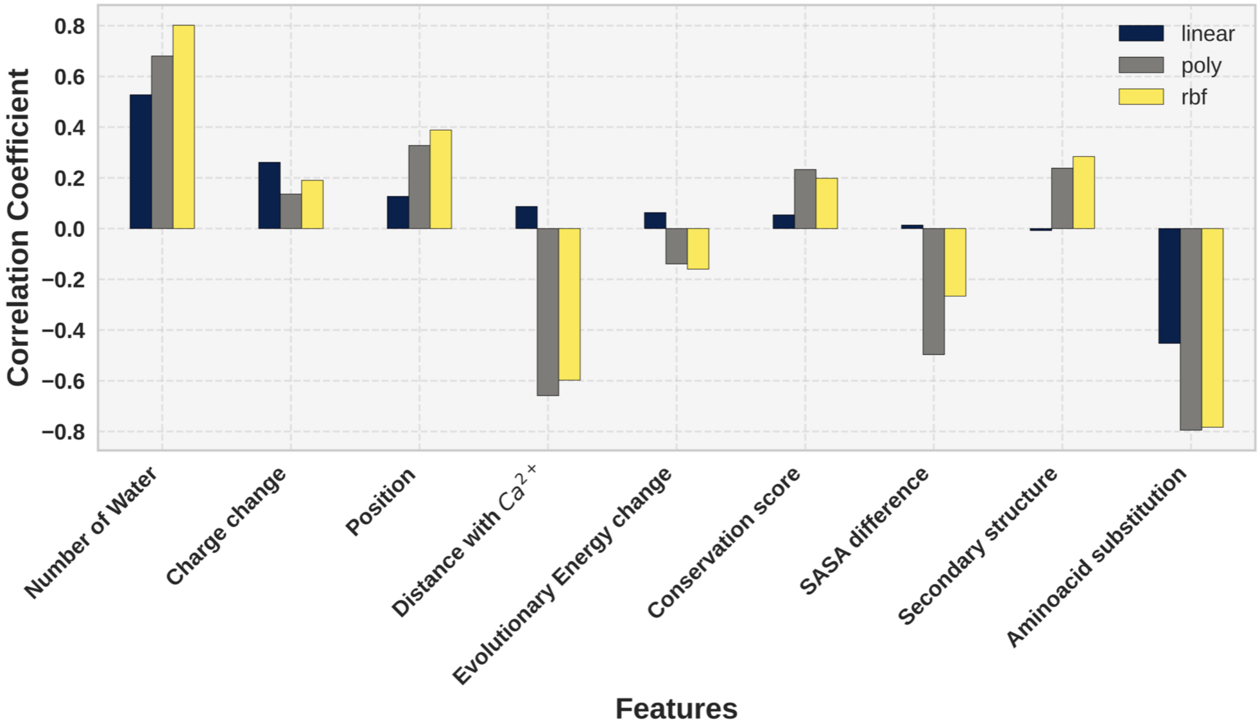


Figure S6: **The barplot graph represents the correlation coefficients of each feature using linear, polynomial, and rbf kernel in support vector regression model depicted by black, grey, and yellow color respectively. The x-axis represents the features used in the the model.**

**Table S1: The partial charges assigned to the $Ca^{2+}$ coordinating and non-coordinating oxygen atoms of cardiac troponin-C protein EF-hand loop residues used in this study.**

| **Charged atomic ions** | **charge(e)** | |
|---|---|---|
| | **ff14SB** | **scaled** |
| ASP 65,67 /O1/O2 | -0.8014 | -0.7389 |
| SER 69/O | -0.5679 | -0.5054 |
| THR 71/O | -0.5679 | -0.5054 |
| GLU 76/O1/O2 | -0.8188 | -0.7563 |

**Table S2: The partial charges assigned to the divalent atom $Ca^{2+}$ and monovalent ions $K^+$ and $Cl^-$ used in this study.**

| **Charged ions** | **charge(e)** | |
|---|---|---|
| | **unscaled** | **scaled** |
| $Ca^{2+}$ | 2.0 | 1.50 |
| $K^+$ | 1.0 | 0.75 |
| $Cl^-$ | 1.0 | -0.75 |

**Table S3: The values taken for the solvent accessible surface area[3] for the amino acid residues used in a feature in Machine Learning model.**

| **Amino acid** | **SASA in $\mathring{A}^2$** |
|---|---|
| Ala | 110.2 |
| Arg | 229.0 |
| Asn | 146.4 |
| Asp | 144.1 |
| Cys | 140.4 |
| Gly | 78.7 |
| Gln | 178.6 |
| Glu | 174.7 |
| His | 181.9 |
| Ile | 185.0 |
| Leu | 183.1 |
| Lys | 205.7 |
| Met | 200.1 |
| Pro | 141.9 |
| Phe | 200.7 |
| Ser | 117.2 |
| Thr | 138.7 |
| Trp | 240.5 |
| Tyr | 213.7 |
| Val | 153.7 |

**Table S4: Calculated binding free energy($\Delta G_{bind}$) and its components $\Delta E_{vdW}$, $\Delta E_{el}$, $\Delta G_{PB}$, $\Delta G_{cav}$, $\Delta G_{disp}$ for cardiac troponin c protein diseased mutations. All values are expressed in kcal/mol, with mean of standard error of the mean provided in parentheses.**

| **System** | $\Delta E_{vdW}$ | $\Delta E_{el}$ | $\Delta G_{PB}$ | $\Delta G_{cav}$ | $\Delta G_{disp}$ | $\Delta G_{bind}$ |
|---|---|---|---|---|---|---|
| WT | 16.89(0.54) | -74.03(0.32) | 52.61(0.14) | -3.48(0.00) | 1.59(0.02) | -6.42(0.44) |
| Y5H | 15.23(0.28) | -74.10(0.20) | 54.54(0.10) | -3.46(0.00) | 1.70(0.01) | -6.09(0.22) |
| A8V | 17.74(0.32) | -77.61(0.20) | 54.61(0.12) | -3.48(0.00) | 1.43(0.01) | -7.31(0.27) |
| F20Q | 21.48(0.18) | -82.47(0.07) | 55.03(0.04) | -3.49(0.00) | 1.34(0.00) | -8.10(0.15) |
| A23Q | 21.29(0.52) | -81.65(0.12) | 54.70(0.12) | -3.49(0.00) | 1.33(0.00) | -7.83(0.44) |
| L29Q | 20.12(0.18) | -79.27(0.07) | 54.40(0.04) | -3.49(0.00) | 1.29(0.00) | -6.95(0.15) |
| A31S | 15.43(0.57) | -74.07(0.34) | 53.70(0.15) | -3.47(0.01) | 1.65(0.02) | -6.76(0.45) |
| S37G | 20.33(0.18) | -79.81(0.07) | 55.04(0.04) | -3.49(0.00) | 1.30(0.00) | -6.63(0.16) |
| E40A | 16.19(0.56) | -71.84(0.36) | 52.18(0.17) | -3.46(0.01) | 1.50(0.03) | -5.42(0.45) |
| V44Q | 19.21(0.58) | -78.35(0.29) | 53.43(0.14) | -3.49(0.00) | 1.33(0.01) | -7.87(0.48) |
| M45Q | 18.80(0.54) | -79.11(0.25) | 55.06(0.13) | -3.48(0.00) | 1.33(0.01) | -7.41(0.45) |
| L48Q | 20.15(0.53) | -79.39(0.20) | 54.38(0.12) | -3.49(0.00) | 1.29(0.00) | -7.07(0.46) |
| Q50R | 20.02(0.55) | -77.74(0.22) | 52.26(0.14) | -3.49(0.00) | 1.33(0.01) | -7.62(0.47) |
| L57Q | 10.39(0.47) | -65.03(0.26) | 50.15(0.16) | -3.22(0.02) | 1.98(0.02) | -5.74(0.38) |
| E59D | 14.09(0.48) | -71.52(0.23) | 52.48(0.13) | -3.47(0.01) | 1.85(0.02) | -6.58(0.41) |
| I61Q | 19.91(0.58) | -76.91(0.24) | 52.62(0.13) | -3.49(0.00) | 1.31(0.01) | -6.56(0.49) |
| D67A | 14.19(0.29) | -73.05(0.37) | 55.38(0.26) | -3.19(0.02) | 1.43(0.01) | -5.25(0.21) |
| D73A | 14.55(0.56) | -64.52(0.26) | 46.64(0.14) | -3.45(0.01) | 1.53(0.02) | -5.25(0.43) |
| D73N | 19.01(0.56) | -69.72(0.23) | 47.07(0.13) | -3.49(0.00) | 1.30(0.01) | -5.82(0.46) |
| D75Y | 14.10(0.18) | -72.17(0.15) | 54.69(0.07) | -3.36(0.01) | 1.61(0.01) | -5.13(0.14) |
| V79Q | 19.33(0.17) | -75.25(0.08) | 50.86(0.04) | -3.49(0.00) | 1.31(0.00) | -7.24(0.15) |
| M81Q | 14.62(0.25) | -71.44(0.14) | 51.97(0.07) | -3.45(0.00) | 1.67(0.01) | -6.63(0.19) |
| C84Y | 18.65(0.19) | -78.28(0.09) | 54.39(0.06) | -3.49(0.00) | 1.34(0.00) | -7.39(0.15) |
| D87A | 16.32(0.29) | -71.81(0.16) | 51.69(0.11) | -3.47(0.00) | 1.52(0.01) | -6.26(0.22) |
| D88A | 14.37(0.33) | -74.49(0.21) | 55.65(0.09) | -3.45(0.00) | 1.68(0.02) | -6.24(0.27) |

**Table S5: Calculated relative binding free energy ($\Delta\Delta G_{bind}$) values for cardiac troponin c protein plausible mutations. All values are expressed in kcal/mol.**

| **System** | $\Delta\Delta G_{cal}$ | System | $\Delta\Delta G_{cal}$ | System | $\Delta\Delta G_{cal}$ |
|---|---|---|---|---|---|
| N18A | -0.16 | L29T | -0.40 | D62A | 0.25 |
| N18D | -1.21 | L29V | -0.12 | D62K | -0.04 |
| N18E | -1.57 | G30D | -1.14 | D62N | 0.18 |
| N18K | -0.47 | G30E | -1.0 | D62R | 0.31 |
| N18Q | -0.61 | G30N | -0.69 | E66A | 0.51 |
| N18S | 0.27 | E32G | 1.22 | E66L | 2.81 |
| A22E | -1.02 | C35A | 0.94 | E66T | 0.21 |
| D25K | 1.87 | C35E | -2.02 | E66V | 2.88 |
| D25S | 0.24 | C35K | -0.89 | V72I | -1.18 |
| I26L | -0.44 | C35T | -0.64 | V72L | -0.56 |
| I26M | 0.45 | S37T | -0.78 | D73E | -0.24 |
| V28D | -0.77 | K43R | -0.23 | D75E | -0.82 |
| L29A | 0.36 | K43T | 1.95 | V79A | -0.05 |
| L29D | -0.75 | K43V | 0.05 | V79K | 0.31 |
| L29E | -0.46 | M47A | 0.20 | V79L | -0.55 |
| L29G | -0.12 | M47S | -0.29 | V79N | 0.26 |
| L29H | 0.38 | P54A | -0.79 | V79S | 0.62 |
| L29K | -0.44 | P54D | 0.77 | V79T | -1.20 |
| L29R | 1.54 | P54E | 0.34 | | |
| L29S | -0.32 | P54K | -1.90 | | |
